# T Cell Calcium Dynamics Visualized in a Ratiometric tdTomato-GCaMP6f Transgenic Reporter Mouse

**DOI:** 10.1101/203950

**Authors:** Tobias X. Dong, Shivashankar Othy, Amit Jairaman, Jonathan Skupsky, Angel Zavala, Ian Parker, Joseph L. Dynes, Michael D. Cahalan

**Affiliations:** Department of Physiology and Biophysics, University of California, 285 Irvine Hall, Irvine, California 92697, USA.; Department of Medicine, University of California, 285 Irvine Hall, Irvine, California 92697, USA.; Department of Neurobiology & Behavior, University of California, McGaugh Hall, Irvine, California 92697, USA.; Institute for Immunology, University of California, 285 Irvine Hall, Irvine, California 92697, USA.

**Keywords:** T cell motility, Orai1, genetically encoded Ca^2+^ indicator, Ca^2+^ signaling, two-photon microscopy

## Abstract

Calcium is an essential cellular messenger that regulates numerous functions in living organisms. Here we describe development and characterization of “Salsa6f”, a fusion of GCaMP6f and tdTomato optimized for cell tracking while monitoring cytosolic Ca^2+^, and a transgenic Ca^2+^ reporter mouse with Salsa6f floxed and targeted to the Rosa26 locus for expression in specific cell types. Using CD4-Cre-Salsa6f mice, we report normal development and function of T cells expressing Salsa6f and demonstrate Ca^2+^ signaling dynamics during T cell receptor engagement in naïve T cells, helper Th17 T cells and regulatory T cells. Salsa6f expression also revealed functional expression of mechanosensitive Piezo1 channels in T cells. Transgenic expression of Salsa6f enables ratiometric imaging of Ca^2+^ signals in complex tissue environments found in vivo. Deep tissue two-photon imaging of T cells in the steady-state lymph node revealed a highly localized Ca^2+^ signaling behavior (“sparkles”) as cells migrate.

## Introduction

Calcium (Ca^2+^) is an essential second messenger responsible for a wide variety of cellular functions (Berridge, Lipp et al. 2000, Clapham 2007, Berridge 2012). Through the use of synthetic small molecule Ca^2+^ indicators such as fura-2 and fluo-4, imaging studies have greatly expanded our understanding of Ca^2+^ signaling dynamics (Tsien, Pozzan et al. 1982, Grynkiewicz, Poenie et al. 1985). However, such indicators cannot be targeted to specific subcellular compartments or cell populations, and are unsuitable for long-term studies due to leakage out of cells. Moreover, they often do not faithfully report pure cytosolic Ca^2+^ signals due to diffusion into other cellular compartments such as the nucleus. One alternative to overcoming these limitations is with genetically encoded Ca^2+^ indicators (GECIs), first developed two decades ago as FRET-based fluorescence probes (Miyawaki, Llopis et al. 1997, Romoser, Hinkle et al. 1997, Perez Koldenkova and Nagai 2013). Key advantages to GECIs include the capability for genetic targeting to specific cell types or subcellular organelles, measuring local Ca^2+^ levels by direct fusion to a protein of interest, modulation of expression levels by inclusion of an inducible promoter, and long term studies due to continuous expression of the genetic indicator (Miyawaki, Llopis et al. 1997, Perez Koldenkova and Nagai 2013). Despite these inherent advantages, the initial FRET-based GECI probes were not widely used as their performance fell far behind small molecule Ca^2+^ indicators, particularly in Ca^2+^ sensitivity, brightness, and dynamic range. Since then, successive rounds of design and contributions from multiple research groups have resulted in numerous variants of GECIs with high dynamic range and dramatically improved performance (Baird, Zacharias et al. 1999, Nakai, Ohkura et al. 2001, Tian, Hires et al. 2009, Zhao, Araki et al. 2011, Akerboom, Chen et al. 2012, Akerboom, Carreras Calderon et al. 2013, Chen, Wardill et al. 2013). Single fluorescent protein-based GECIs containing a circularly permutated green fluorescent protein (GFP) exhibit high brightness, fast response kinetics, and offer multiple color variants, including the GECO and the GCaMP series (Tian, Hires et al. 2009, Zhao, Araki et al. 2011, Akerboom, Chen et al. 2012, Chen, Wardill et al. 2013). FRET-based GECIs have continued to evolve as well, with sequential improvements including incorporation of circularly permuted yellow fluorescent proteins (cpYFPs) to improve dynamic range in the yellow cameleon (YC) family (Nagai, Yamada et al. 2004), use of troponin C as the Ca^2+^ sensing element in the TN indicator family (Heim and Griesbeck 2004), computational redesign of the calmodulin-M13 interface to increase the range of Ca^2+^ sensitivity and reduce perturbation by native calmodulin in the DcpV family (Palmer, Giacomello et al. 2006), and complete redesign of the troponin C domain to increase response kinetics and reduce buffering of cytosolic Ca^2+^ in the TN-XXL family (Mank, Reiff et al. 2006, Mank, Santos et al. 2008).

The latest generation of GECIs have crossed key performance thresholds previously set by small-molecule indicators, enabling GECIs to be widely applied in diverse Ca^2+^ imaging studies without sacrificing performance. Members of the GCaMP6 family are capable of tracking cytosolic Ca^2+^ changes from single neuronal action potentials, with higher sensitivity than small-molecule indicators such as OGB-1 (Chen, Wardill et al. 2013). The availability of multicolored variants in the GECO family and the RCaMP series allowed for simultaneous measurement of Ca^2+^ dynamics in different cell populations in the same preparation, or in different subcellular compartments within the same cell (Zhao, Araki et al. 2011, Akerboom, Carreras Calderon et al. 2013). These variants can be integrated with optogenetics to simultaneously evoke channel rhodopsin activity while monitoring localized Ca^2+^ responses in independent spectral channels (Akerboom, Carreras Calderon et al. 2013). Moreover, individual GECIs can be tagged onto membrane Ca^2+^ channels to directly measure Ca^2+^ influx through the target channel of interest, enabling optical recording of single channel activity without the need for technique-intensive patch clamping (Dynes, Amcheslavsky et al. 2016).

Another advantage of GECIs is the capability to be incorporated into transgenic organisms. Although several GECI-expressing transgenic mouse lines have already been reported, many of these studies used older variants of GECIs that are expressed only in selected tissues (Hasan, Friedrich et al. 2004, Ji, Feldman et al. 2004, Tallini, Ohkura et al. 2006, Heim, Garaschuk et al. 2007). The Ai38 mouse line overcomes these issues by combining GCaMP3 with a robust and flexible Cre/lox system for selective expression in specific cell populations (Zariwala, Borghuis et al. 2012). Based on a series of Cre-responder lines designed for characterization of the whole mouse brain (Madisen, Zwingman et al. 2010), the Ai38 mouse line contains GCaMP3 targeted to the Rosa26 locus but requires Cre recombinase for expression. By crossing Ai38 with various Cre mouse lines, GCaMP3 can be selectively expressed in specific cell populations. Thus, target cells may be endogenously labeled without invasive procedures, avoiding potential off-target side effects reported in GECI transgenic lines with global expression (Direnberger, Mues et al. 2012). The newly released PC::G5-tdT mouse line provides improved functionality by targeting a Cre-dependent GCaMP5G-IRES-tdTomato transgenic cassette to the *Polr2a* locus (Gee, Smith et al. 2014). However, in the PC::G5-tdT mouse line, GCaMP5G and tdTomato are expressed individually, and localize to different cell compartments. Since expression of tdTomato is driven by an internal ribosomal entry site, the expression level is highly variable and weaker than GCaMP5G, limiting identification of positive cells and preventing accurate ratiometric measurements.

Although single fluorescent protein-based indicators have high brightness and fast response kinetics, as non-ratiometric probes they are problematic for Ca^2+^ imaging in motile cells where fluorescence changes resulting from movement are indistinguishable from actual changes in Ca^2+^ levels. Here, we introduce a novel genetically encoded Ca^2+^ indicator - that we christen ‘Salsa6f’ - by fusing green GCaMP6f to the Ca^2+^-insensitive red fluorescent protein tdTomato. This probe enables true ratiometric imaging, in conjunction with the high dynamic range of GCaMP6. We further describe the generation of a transgenic mouse enabling Salsa6f expression in a tissue-specific manner, and demonstrate its utility for imaging cells of T lymphocytes in vitro and in vivo.

## Results

### A novel ratiometric genetically encoded Ca^2+^ indicator, Salsa6f

In order to develop a better tool to monitor Ca^2+^ signaling in T cells both in vivo and in vitro, we first evaluated the latest generation of genetically encoded Ca^2+^ indicators (GECIs) (Zhao, Araki et al. 2011, Chen, Wardill et al. 2013). A variety of single fluorescent protein-based GECIs were transiently expressed and screened in HEK 293A cells (**Figure 1A**), and GCaMP6f was selected based on fluorescence intensity, dynamic range, and Ca^2+^ affinity suitable for detecting a spectrum of cytosolic Ca^2+^ signals (*K_d_* = 375 nM). To enable cell tracking even when basal Ca^2+^ levels evoke little GCaMP6f fluorescence, we fused GCaMP6f to the Ca^2+^-insensitive red fluorescent protein tdTomato, chosen for its photostability and efficient two-photon excitation (Drobizhev, Makarov et al. 2011). A V5 epitope tag (Lobbestael E 2010) serves to link tdTomato to GCaMP6f (**Figure 1C**). The resultant ratiometric fusion indicator, coined “Salsa6f” for the combination of red tdTomato with the green GCaMP6f, was readily expressed by transfection into HEK 293A cells and human T cells. Salsa6f exhibited a ten-fold dynamic range, with a brightness comparable to GCaMP6f alone (**Figure 1A,B**). For two-photon microscopy, both components of Salsa6f can be visualized by femtosecond excitation at 900 nm (**Figure 1D**). GCaMP6f produces increased green fluorescence during elevations in cytosolic Ca^2+^, while tdTomato provides a stable red fluorescence that facilitates cell tracking and allows for ratiometric Ca^2+^ imaging (**Figure 1D; Video 1**). Salsa6f is excluded from the nucleus, ensuring accurate measurement of cytosolic Ca^2+^ fluctuations (**Figure 1D,E**). When expressed by transfection in human T cells, Salsa6f reported Ca^2+^ oscillations induced by immobilized αCD3/28 antibodies with a high signal to noise ratio and time resolution (**Figure 1E,F**).

**Figure 1.**
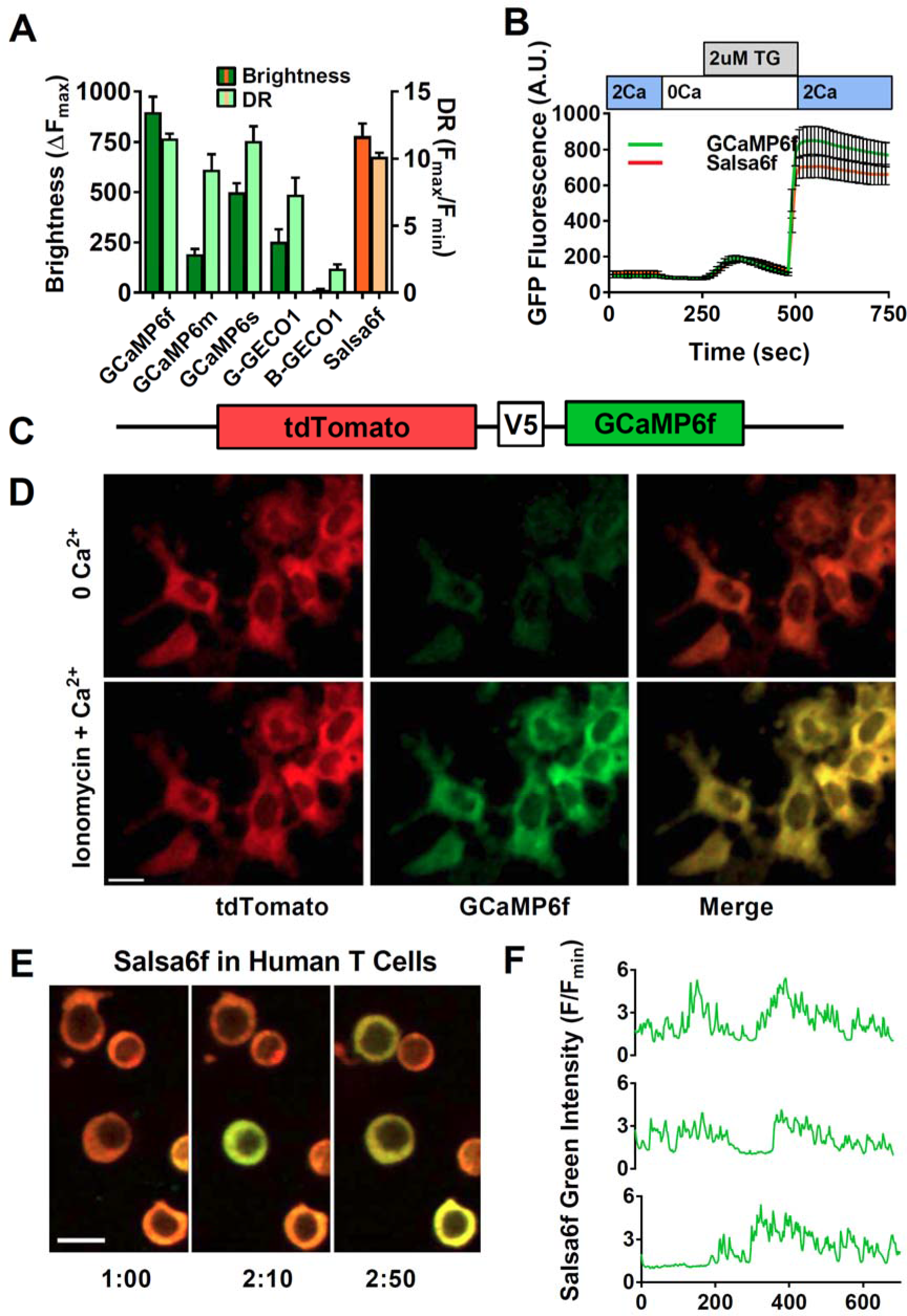
Design of novel tdTomato-V5-GCaMP6f fusion probe “Salsa6f” and characterization in living cells. (**A**) Several genetically encoded Ca^2+^ indicators were screened in vitro in HEK 293A cells, by co-transfecting with Orai1/STIM1 and measuring Ca^2+^ influx after thapsigargin-induced store depletion, showing maximum change in fluorescence intensity in dark green bars and dynamic range (DR) in light green bars, with Salsa6f shown in orange bars on right; n > 30 cells per probe, from two different transfections, error bars indicate SEM. (**B**) Averaged thapsigargin-induced Ca^2+^ entry, measured by change in GFP fluorescence, in GCaMP6f (green, 11.5 ± 0.3, n = 63) or Salsa6f (orange, 10.2 ± 0.3, n = 78) transfected HEK cells; data from two different transfections, error bars indicate SEM. (**C**) Diagram of Salsa6f construct used in transfection. (**D**) Two-photon images of Salsa6f co-transfected in HEK cells with Orai1/STIM1, showing red (tdTomato), green (GCaMP6f), and merged channels, at baseline in 0 mM extracellular Ca^2+^ and after maximum stimulation with 2 μM ionomycin in 2 mM extracellular Ca^2+^; scale bar = 20 μm; see **Video 1**; data representative of at least three different experiments. (**E**) Confocal time lapse microscopy of human CD4^+^ T cells transfected with Salsa6f, then activated for two days on plate-bound αCD3/28 antibodies; time = min:sec, scale bar = 10 μm. (**F**) Representative cell traces of activated human T cells transfected with Salsa6f, tracking green fluorescence intensity only; data representative of at least three different experiments.

### Generation of Salsa6f transgenic reporter mice and validation in immune cells

Guided by the transgenic targeting strategy for the Ai38 mouse line (Zariwala, Borghuis et al. 2012), we inserted Salsa6f into a *ROSA26*-pCAG-LSL-Salsa6f-WPRE-bGHpA-NeoR cassette, then targeted it to the Rosa26 locus in JM8.N4 mouse embryonic stem (ES) cells (**Figure 2A**). Insertion events were selected by neomycin resistance, and correctly targeted clones were screened by Southern blot (**Figure 2B**), then injected into C57BL/6J blastocysts for implantation. Chimeric pups carrying the Salsa6f transgene were identified by PCR screening for the *Nnt* gene, as the initial JM8.N4 ES cells were *Nnt*^+/+^ while the C57BL/6J blastocysts were *Nnt*^−/−^ (**Figure 2C**). Positive chimeras were bred to R26ΦC31o mice to remove the neomycin resistance gene and to produce Salsa6f^LSL/−^ F1 founders, then further bred to generate homozygotic Salsa6f^LSL/LSL^ mice.

**Figure 2.**
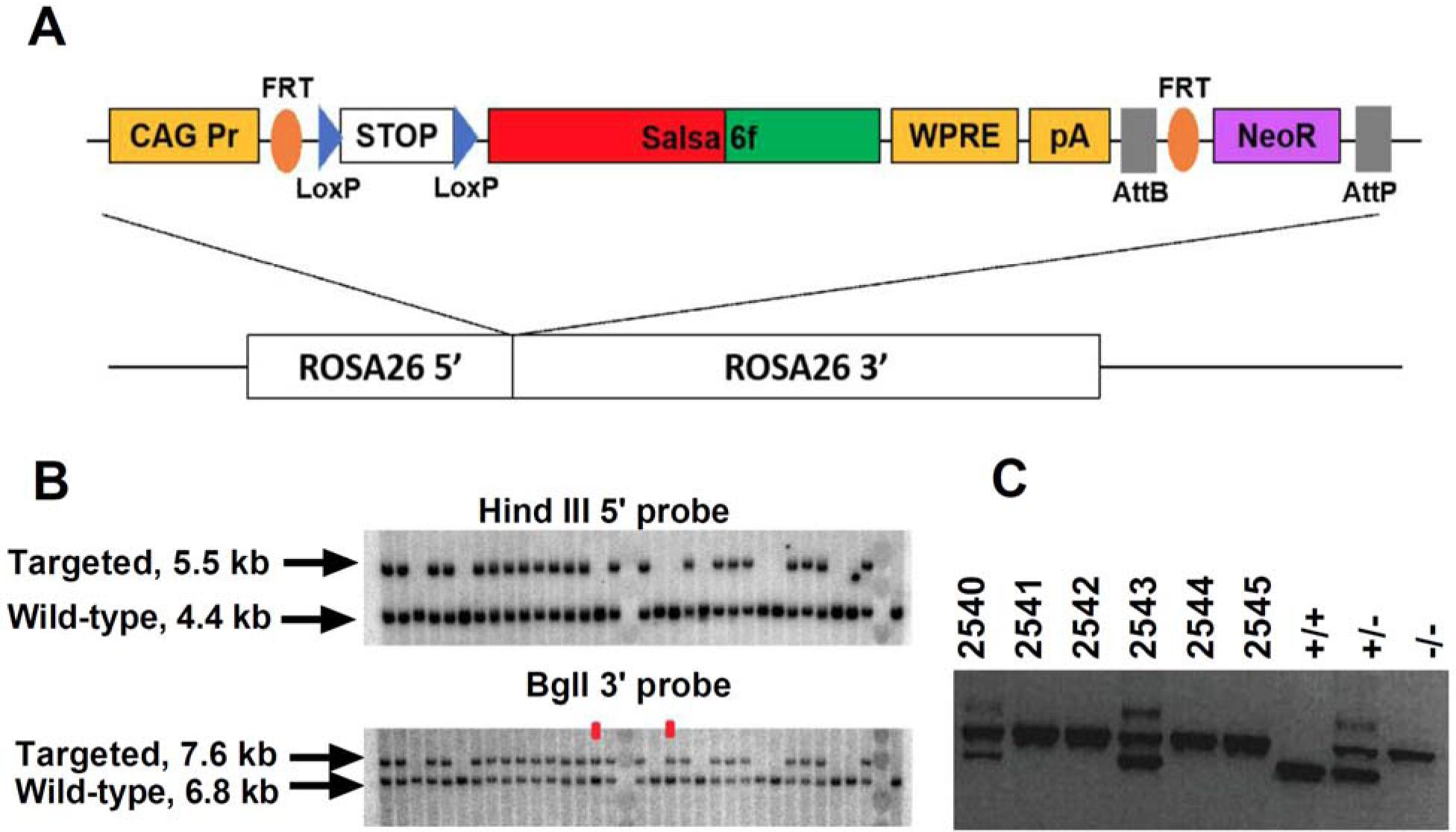
Generation of Salsa6f transgenic mouse line targeted to Rosa26 locus. (**A**) Transgenic targeting vector for Salsa6f, inserted between Rosa26 homology arms and electroporated into embryonic stem cells. CAG Pr: cytomegalovirus early enhancer/chicken β-actin promoter; Salsa6f: tdTomato-V5-GCaMP6f; FRT, LoxP, AttB, AttP: recombinase sites; WPRE: woodchuck hepatitis virus posttranscriptional regulatory element; pA: bovine growth hormone polyadenylation sequence; NeoR: neomycin resistance gene. (**B**) Correctly targeted ES cells were screened by Southern blot after HindIII digest for the 5’ end (top) or BglI digest for the 3’ end (bottom). The two clones marked in red failed to integrate at the 5’ end. (**C**) PCR screening for chimeras based on presence of the Nnt mutation, present only in JM8.N4 ES cells but not in the C57BL/6J blastocyst donors. 2540 and 2543 are chimeras. Control lanes on the right are wildtype (*Nnt*^+/+^), heterozygous (*Nnt*^+/−^), or homozygous mutant (*Nnt*^−/−^).

Salsa6f^LSL/LSL^ mice were bred to CD4-Cre^+/+^ mice to obtain CD4-Cre^+/−^ Salsa6f^+/−^ reporter mice, designated as CD4-Salsa6f^+/−^ mice from here on, that selectively express Salsa6f in T cells (**Figure 3A**). Salsa6f was detected by tdTomato fluorescence on flow cytometry. 88% of Salsa6f^+^ cells in thymus were double positive for CD4 and CD8 (**Figure 3B**). This is due to the double-positive stage during development, in which developing thymocytes will express both CD4 and CD8 before undergoing positive and negative selection to become either mature CD4^+^ or CD8^+^ T cells. Salsa6f was readily detected in cells from spleen (40%), lymph node (57%), and thymus (93%) (**Figure 3C**). As expected, double positive cells were not detected in the spleen (**Figure 3D**). More than 98% of CD4^+^ and CD8^+^ T cells from these reporter mice were positive for Salsa6f. Salsa6f was also detected in 5% of CD19^+^ cells and 3% of CD11b^+^ cells (**Figure 3E**). A small fraction of B cells express CD4 mRNA, which may explain the presence of Salsa6f in CD19^+^ cells (Zhang and Henderson 1994). CD11b^+^ cells positive for Salsa6f may be splenic resident dendritic cells that also express CD4 (Vremec, Pooley et al. 2000, Turley, Fletcher et al. 2010). The total number and relative frequencies of CD4^+^, CD8^+^, CD19^+^, and CD11b^+^ cells were similar to the CD4-Cre controls (**Figure 3F,G**).

**Figure 3.**
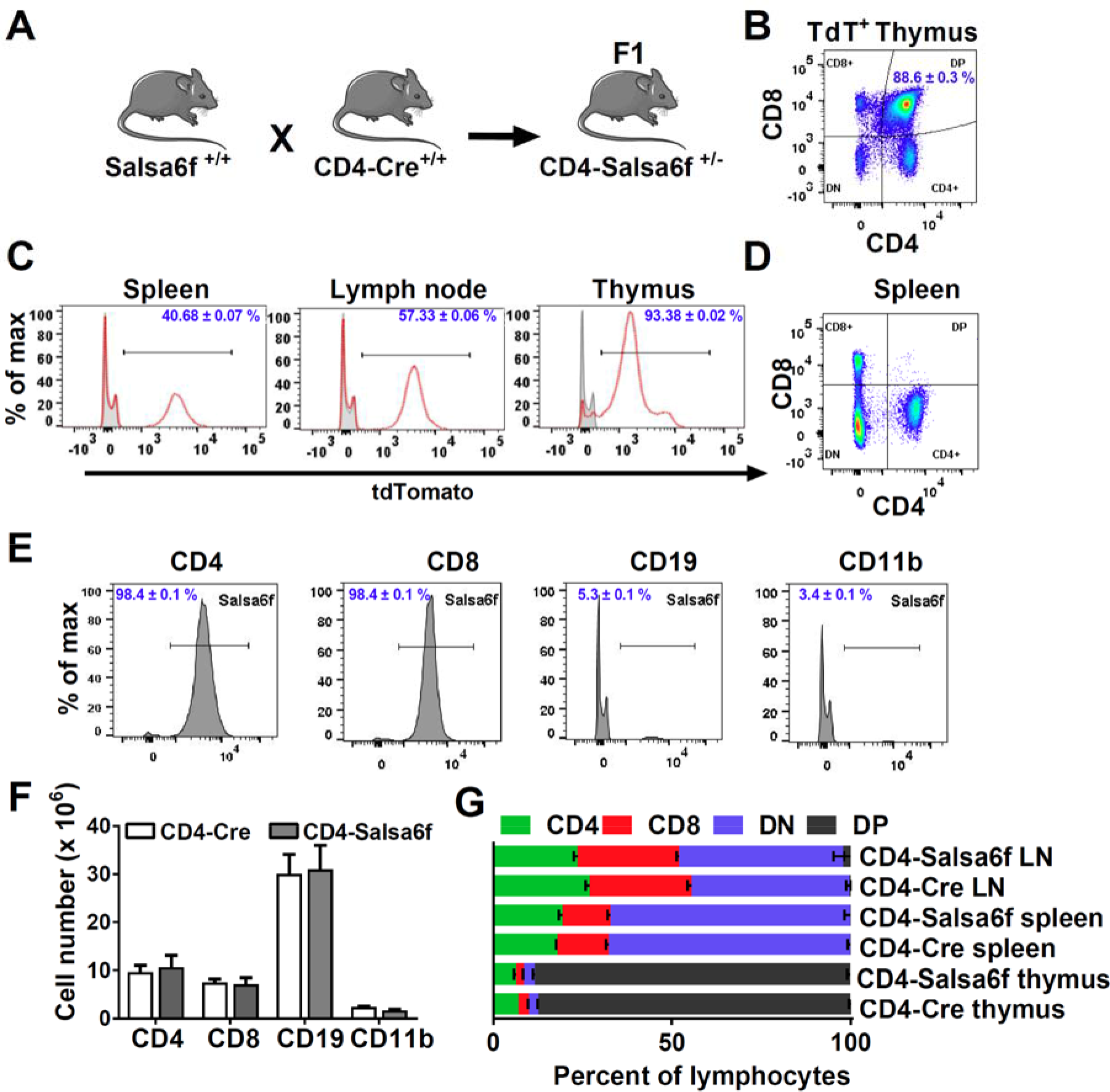
CD4-Salsa6f mice show normal immune cell development and expression. (**A**) Experimental design to target expression of Salsa6f in CD4 cells. (**B**) CD4, CD8 and double-positive cells gated on tdTomato (Salsa6f^+^ cells) from thymus. (**C**) Histograms showing percent of Salsa6f^+^ cells in spleen, LN, and thymus. (**D**) CD4, CD8, and double positive cells from spleen, gated on tdTomato (Salsa6f^+^ cells). (**E**) Histograms showing percent of Salsa6f^+^ cells within CD4, CD8, CD19, CD11b populations from spleen. (**F**) Total number of CD4, CD8, CD19, CD11b cells in the spleen of CD4-Salsa6f^+/−^ mice and CD4-Cre mice (n=6 mice). (**G**) Relative percentages of CD4, CD8, CD19, CD11b cells in thymus, lymph nodes, and spleen of CD4-Salsa6f mice and CD4-Cre mice (n=6).

To evaluate functional responses downstream of Ca^2+^ signaling in Salsa6f-expressing T cells, we first purified CD4^+^ T cells and monitored cell proliferation in vitro during TCR engagement of αCD3 and co-stimulating αCD28 antibodies attached to activating beads. Salsa6f-expressing CD4^+^ T cells proliferated similar to the CD4-Cre controls (**Figure 4A,B**). To further probe functional responses, we differentiated naive CD4^+^ T cells during polarizing cytokine stimuli to generate Th1, Th17 and induced regulatory T cells (iTregs). Salsa6f^+^ naive CD4^+^ T cells readily differentiated into various helper T cell subtypes similar to the CD4-Cre controls (**Figure 4C-E**). In addition, as described in the companion paper, adoptively transferred Salsa6f^+^ cells readily homed to lymph nodes where they exhibited normal motility. In summary, our results demonstrate normal T cell function of CD4-Salsa6f^+/−^ T cells with respect to cellular phenotype, cell proliferation, differentiation, homing, and motility.

**Figure 4.**
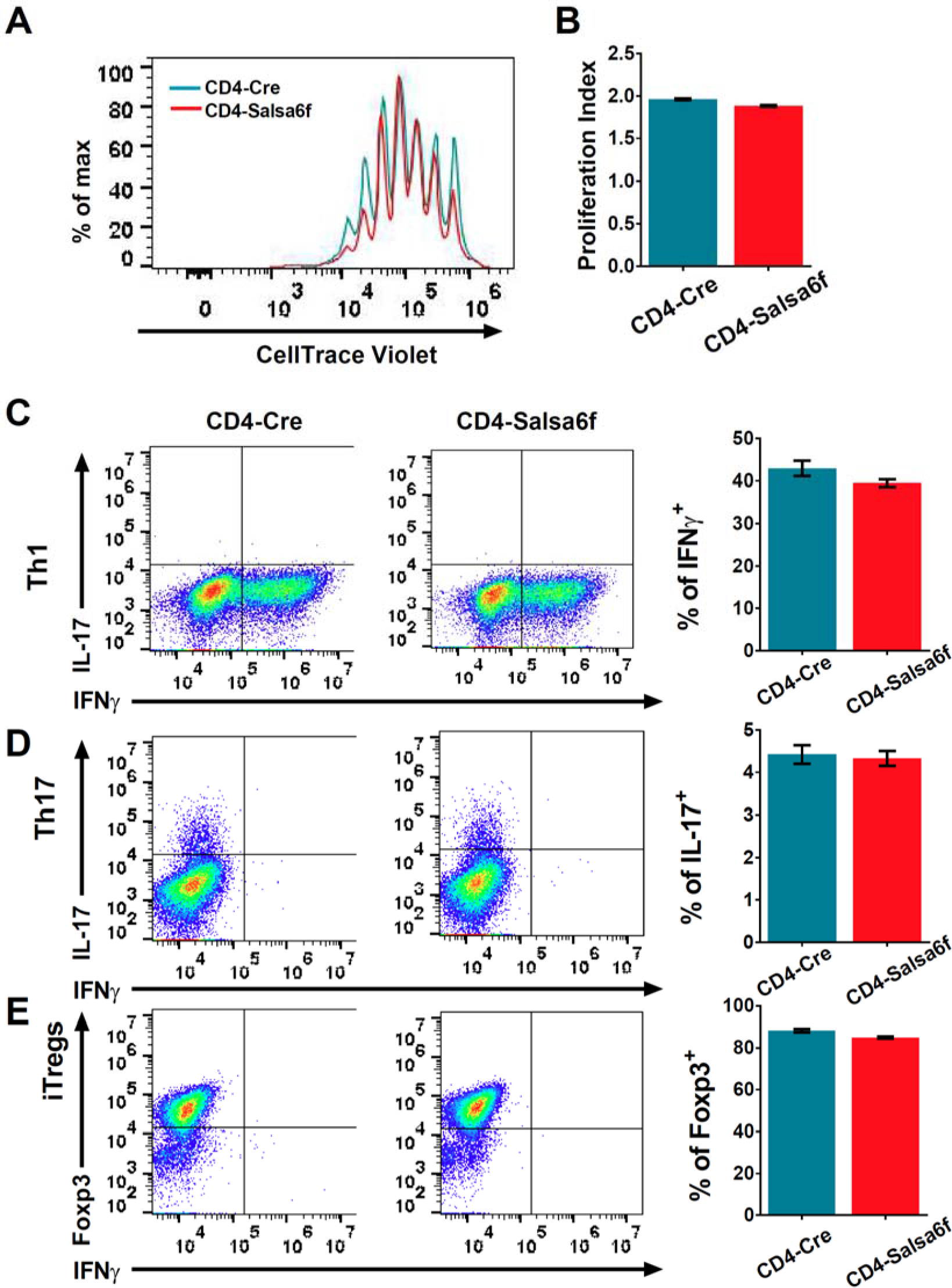
Functional responses of CD4-Salsa6f T cells in vitro. (**A**) Representative histogram showing cell trace violet (CTV) dilution in CD4-Cre (teal) and CD4-Salsa6f^+/−^ T cells (red) at 92 hours following stimulation with αCD3/28 Dynabeads (1:1 ratio). (**B**) Proliferation index measured on CTV dilution curves (n=10). (**C-E**) Dot plots showing differentiation of naïve T cells from CD4-Cre and CD4-Salsa6f^+/−^ mice into Th1 cells (**C**), Th17 cells (**D**) and iTregs (**E**) after 6 days (n = 4 mice). Right panels show average percent of IFNγ+ cells (**C**), IL-17^+^ cells (**D**) and Foxp3^+^ cells (**E**).

### Single-cell ratiometric Ca^2+^ measurement in CD4-Salsa6f reporter mice

CD4^+^ T cells were purified from CD4-Salsa6f^+/−^ reporter mice, stimulated with plate-bound αCD3/28 antibodies for two days, and imaged by confocal microscopy while still in contact with immobilized antibodies. Red and green fluorescence emitted from the cytosol of individual cells was tracked (**Figure 5A**, **Video 2**). Activated CD4^+^ T cells expressing Salsa6f exhibited stable red fluorescence and wide fluctuations in green fluorescence due to Ca^2+^ oscillations resulting from T cell receptor engagement (**Figure 5B**). Despite variability in total fluorescence between cells due to individual differences in cell size, the basal and peak green/red Salsa6f ratios (referred from now on as G/R ratio for GCaMP6f/tdTomato intensity) were comparable between cells and showed up to six-fold increases during peaks in Ca^2+^ fluctuations. This level of response matches our previous experiments in activated human T cells transfected with Salsa6f (*c.f.*, **Figure 1E,F**), and supports the consistency in making ratiometric measurements with Salsa6f. Flow cytometric analysis of Salsa6f^+/−^ mouse T cells revealed a thirteen-fold increase in G/R ratio, by pretreatment with ionomycin in free Ca^2+^ to deplete cytosolic Ca^2+^ followed by addback of extracellular Ca^2+^, further emphasizing the high dynamic range of Salsa6f (**Figure 5D**). Finally, to test if increasing the genetic dosage can improve the brightness of Salsa6f, we compared CD4^+^ T cells from heterozygotic CD4-Salsa6f^+/−^ mice and homozygotic CD4-Salsa6f^+/+^ mice. T cells from homozygous mice with two allelic copies of the Salsa6f reporter cassette exhibited almost a two-fold increase in tdTomato fluorescence compared to heterozygous mice (**Figure 5E**), allowing for genetic control of Salsa6f expression level when brightness is an issue.

**Figure 5.**
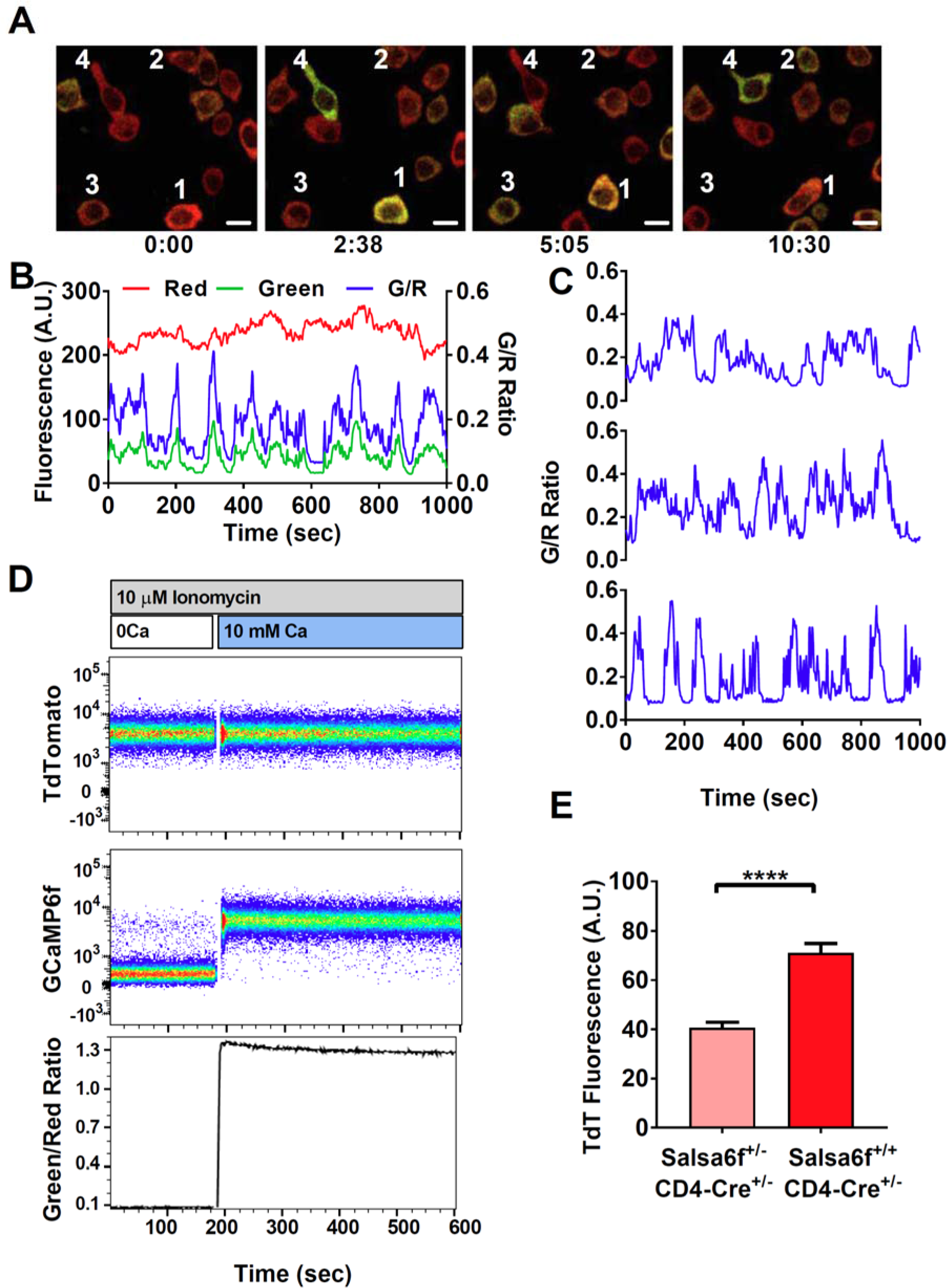
Single-cell readout of Salsa6f calcium signals in T cells. (**A**) Confocal imaging of Ca^2+^ signals in activating CD4^+^ T cells from CD4-Salsa6f^+/−^ mice, after two day stimulation on plate bound αCD3/28 antibody, showing merged green (GCaMP6f) and red (tdTomato) channels; time = min:sec; scale bar = 10 μm. (**B**) Representative traces from cell #3 in (**A**), showing total fluorescence intensity changes in GCaMP6f (green), tdTomato (red), and green/red ratio (G/R, blue). (**C**) G/R ratios for cells 1, 2, and 4 from (**A**). (**D**) Dynamic range of Salsa6f in resting CD4 T cells, measured as green/red fluorescence by flow cytometry. Cells were pre-treated with 10 μM ionomycin in Ca^2+^-free solution (white bar), followed by re-addition of 10 mM Ca^2+^ (blue bar). (**E**) Averaged tdTomato fluorescence in resting T cells from heterozygous CD4-Salsa6f^+/−^ compared to homozygotic CD4-Salsa6f^+/+^ mice.

### Cytosolic localization and calibration of Salsa6f in transgenic T lymphocytes

We first examined the localization of Salsa6f in naïve CD4^+^ T cells isolated from CD4-Salsa6f^+/−^ mice and in CD4^+^ T cells activated for 2 days on plate-bound αCD3/28. Line scans of the confocal images of cells plated on poly-L-lysine coated coverslips showed that Salsa6f is primarily localized to the cytoplasm and is excluded from the nucleus (**Figure 6A,B**). Increasing the cytosolic Ca^2+^ levels using thapsigargin (TG) in 2 mM Ca^2+^ Ringer’s solution caused a selective increase in the GCaMP6f signal, without altering the localization of Salsa6f probe. In contrast, the chemical Ca^2+^ indicators fluo-4 or fura-2 loaded into CD4^+^ T cells from CD4-Cre mice are localized throughout the cell, including the nucleus (**Figure 6C** and data not shown). A different transgenic mouse, CD4-Cre 5GtdT^+/−^, utilizes an internal ribosomal entry site to express both tdTomato and GCaMP5G as separate proteins that localize differently in cells (Gee, Smith et al. 2014), tdTomato throughout the cell including the nucleus and GCaMP5G predominantly in the cytosol (**Figure 6-figure supplement 1**). In contrast, our tandem probe, Salsa6f, results in both red and green fluorescent proteins co-localized in the cytosol, allowing true ratiometric Ca^2+^ imaging and facilitating tracking of cells.

**Figure 6.**
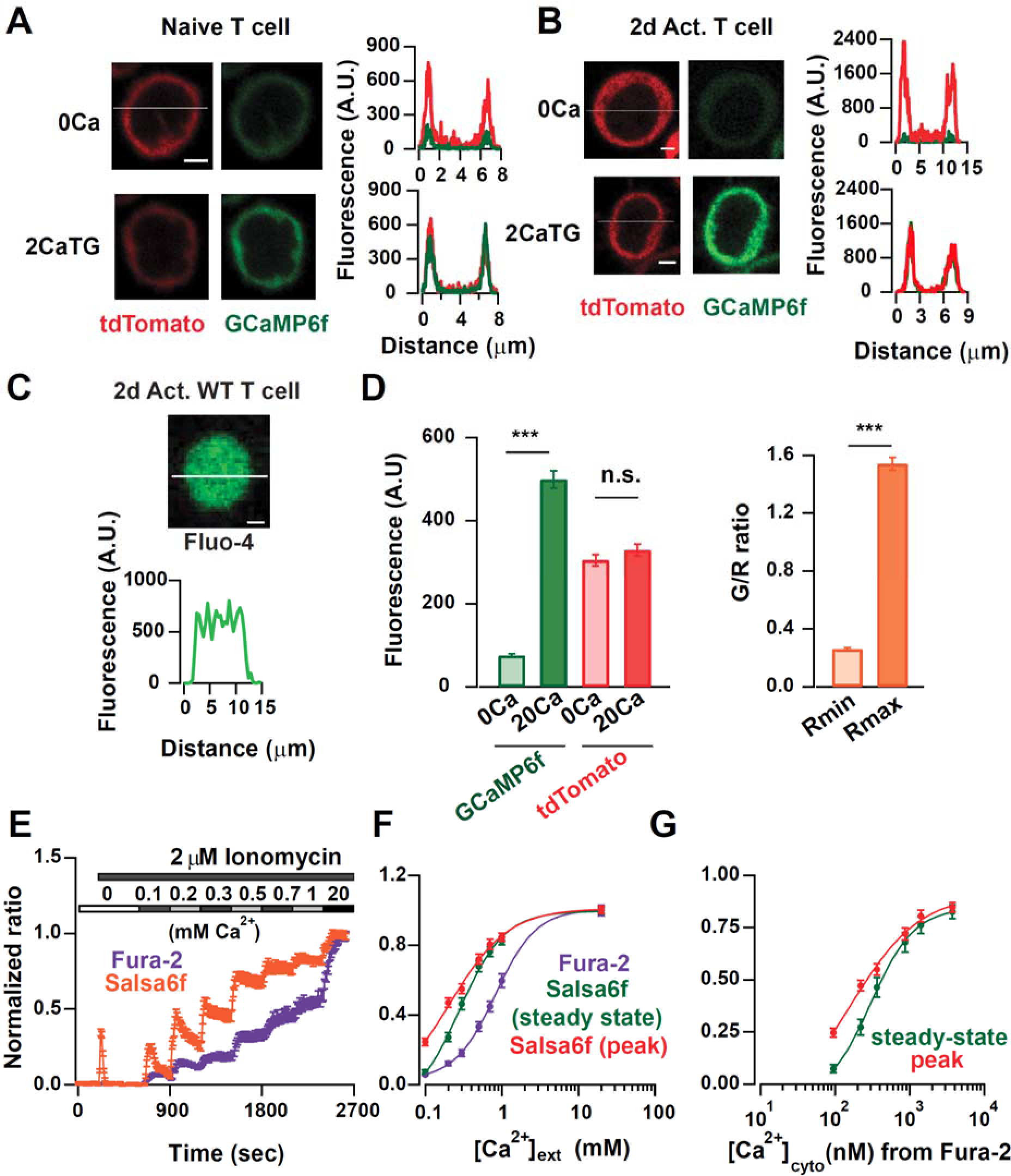
Probe characterization and calibration of [Ca^2+^] in Salsa6f T cells. (**A**) Confocal image of a naïve T cell from a CD4-Salsa6f^+/−^ mouse. Upper panel: tdTomato (left) and GCaMP6f (right) fluorescence intensity in Ca^2+^-free Ringer solution. Lower panel: same cell treated with 2 μM thapsigargin (TG) in Ringer solution containing 2 mM Ca^2+^. Line scan for each condition is shown adjacent to the images. Scale bar = 2 μ.m for **A-C**. (**B)** Confocal images of Salsa6f localization in a 2-day activated CD4^+^ T-cell from CD4-Salsa6f^+/−^ mouse. **(C)** Confocal image of a Fluo-4 (5 μM) loaded CD4^+^ T cell from CD4-Cre mouse. **(D)** Average GCaMP6f and tdTomato intensities and G/R ratios in 2-day activated CD4^+^ T cells treated with 2 μM ionomycin in Ca^2+^ free buffer (F_min_) and 20 mM Ca^2+^ buffer (F_max_); n = 76 cells, representative of 3 experiments. **(E)** Average 340 / 380 nm ratios in fura-2 loaded CD4^+^ T cells (n=59 cells) and G/R ratios in Salsa6f CD4^+^ T cells (n=47 cells) treated identically with 2 μM ionomycin followed by graded increases of external Ca^2+^ concentration as indicated. **(F)** Steady-state fura-2 and Salsa6f ratios recorded 300 s after solution application and peak Salsa6f ratio from **6E** plotted as a function of external Ca^2+^ concentration. **(G)** Steady-state and peak Salsa6f ratios plotted as a function of cytosolic Ca^2+^ concentrations calculated from the fura-2 experiment, assuming a fura-2 *K_d_* of 225 nM. The points were fit with a 4 parameter Hill equation to obtain the *K_d_* for Salsa6f, with the following parameters: Salsa6f steady-state: Hill coefficient = 1.49 ± 0.16; *K_d_* = 301 ± 24; Salsa6f peak: Hill coefficient = 0.93 ± 0.4; *K_d_* = 162 ± 48. Data are representative of three experiments.

To estimate the dynamic range of the probe in Salsa6f-expressing T cells, we treated the cells with ionomycin in Ca^2+^-free and 20 mM Ca^2+^ Ringer’s solution to get the minimum and maximum fluorescence signals and ratios (F_max_, R_max_, F_min_ and R_min_). Addition of high Ca^2+^ buffer resulted in a robust and selective increase in the GCaMP6f signal without significantly affecting the tdTomato signal (**Figure 6D**). Based on the fold change in the G/R ratio, the dynamic range of the probe was calculated to be ~ 5.5-fold.

GCaMP6f is reported to have a *K*_*d*_ of 290 nM at 25° C and 375 nM at 37° C based on in vitro measurements with purified protein (Chen, Wardill et al. 2013, Badura, Sun et al. 2014). In situ *K*_*d*_ values of Ca^2+^ indicators often vary significantly from the reported in vitro values (Negulescu and Machen 1990), although to our knowledge, the in situ *K*_*d*_ of GCaMP6f has not been determined in any specific cell type. To characterize the Ca^2+^ affinity and binding kinetics of Salsa6f in situ, we performed time-lapse imaging at 25° C on 2 day-activated CD4^+^ cells isolated from CD4-Salsa6f^+/−^ mice and plated on poly-L-lysine. We recorded G/R ratios in response to ionomycin and applied stepwise increases in the external Ca^2+^ concentration (**Figure 6E**). Our strategy for calibration was to compare these results with Ca^2+^ signals in fura-2-loaded CD4^+^ T cells from CD4-Cre mice using exactly the same protocol of cell isolation, plating, and solution application (**Figure 6E,F**), the rationale being that fura-2 has an in situ *K_d_* of around 225 nM at 25° C (Lewis and Cahalan 1989), which is close to the range of in vitro *K*_*d*_ values reported for GCaMP6f (Chen, Wardill et al. 2013, Badura, Sun et al. 2014). For meaningful comparison, the traces were normalized with R_min_=0 and R_max_=1 for both fura-2 and Salsa6f. To our surprise, Salsa6f responded with faster rise times and at lower external Ca^2+^ concentrations than did fura-2 to progressive increases in cytosolic Ca^2+^ levels. These observations were unexpected, given that the reported in situ Ca^2+^ affinity of fura-2 is higher than the in-vitro affinity of GCaMP6f. Additionally, the Ca^2+^ signal measured with Salsa6f decayed faster than that measured with fura-2. This effect was more prominent at lower cytosolic Ca^2+^ levels and as the Ca^2+^ levels increased, the rate of decay diminished and the overall kinetics were then similar to fura-2. The faster rise and fall times of the Ca^2+^ signals seen in Salsa6f expressing cells is not likely to be due to differential activity of Ca^2+^ influx and efflux mechanisms between CD4-Cre and CD4-Salsa6f^+/−^ cells since fura-2 signals in Salsa6f cells also displayed kinetics very similar to that seen in WT CD4-Cre cells (data not shown).

Secondly, Salsa6f responses saturated at lower cytosolic Ca^2+^ levels than fura-2 responses. This is not altogether surprising given that genetically encoded Ca^2+^ indicators have been reported to have a steeper Hill coefficient than chemical indicators (Badura, Sun et al. 2014). Obtaining steady-state cytosolic Ca^2+^ concentrations from fura-2 measurements in WT CD4-Cre cells, and assuming that CD4-Salsa6f^+/−^ cells reach similar Ca^2+^ levels, we plotted both the peak and the steady state Salsa6f G/R ratios against the cytosolic Ca^2+^ concentrations obtained from the fura-2 experiment (**Figure 6G**). Using steady-state levels for Salsa6f, we calculated a *K*_*d*_ of ~300 nM, while using peak levels for Salsa6f probe gave a *K*_*d*_ of ~160 nM. Furthermore, Salsa6f was sensitive in detecting cytosolic Ca^2+^ in the range of 100 nM - 2 μM. Based on these results, we conclude that Salsa6f probe with its high sensitivity is well suited to detect small changes in cytosolic Ca^2+^ in response to various physiological stimuli, while its excellent dynamic range also allows us to detect larger elevations in Ca^2+^ up to 2 μM.

### T cell Ca^2+^ signaling in response to Ca^2+^ store depletion, T cell receptor engagement, and mechanical stimulation

TCR engagement activates a canonical Ca^2+^ signaling pathway in T cells, characterized by IP_3_-induced Ca^2+^ release from the endoplasmic reticulum, leading to store-operated Ca^2+^ entry (SOCE) through Orai1 channels (Cahalan and Chandy 2009, Prakriya and Lewis 2015). Past studies on T cell Ca^2+^ signaling have largely relied on chemical indicators like fura-2 and fluo-4 which have the drawback of being distributed into nucleus and other cellular compartments, thus confounding the measurement of pure cytoplasmic Ca^2+^ signals, a problem particularly notable in T cells with its large nuclear to cytoplasmic volume. Salsa6f, with its large dynamic range, ratiometric readout and targeted localization in the cytosol, is thus well suited to record physiological Ca^2+^ signals that are primarily cytosolic. To this end, we recorded Ca^2+^ signals from 2 day-activated CD4^+^ T cells from CD4-Salsa6f^+/−^ mice in response to a variety of stimuli.

To study SOCE more directly, we depleted ER Ca^2+^ stores with TG in Ca^2+^ free solution. We observed a small but sharp initial peak indicating ER store release and a sustained Ca^2+^ signal upon restoring Ca^2+^ to the external bath suggestive of SOCE (**Figure 7A, Video 3**). Single traces revealed that almost all cells responded to this supra-physiological stimulus (**Figure 7A**, right panel).

**Figure 7.**
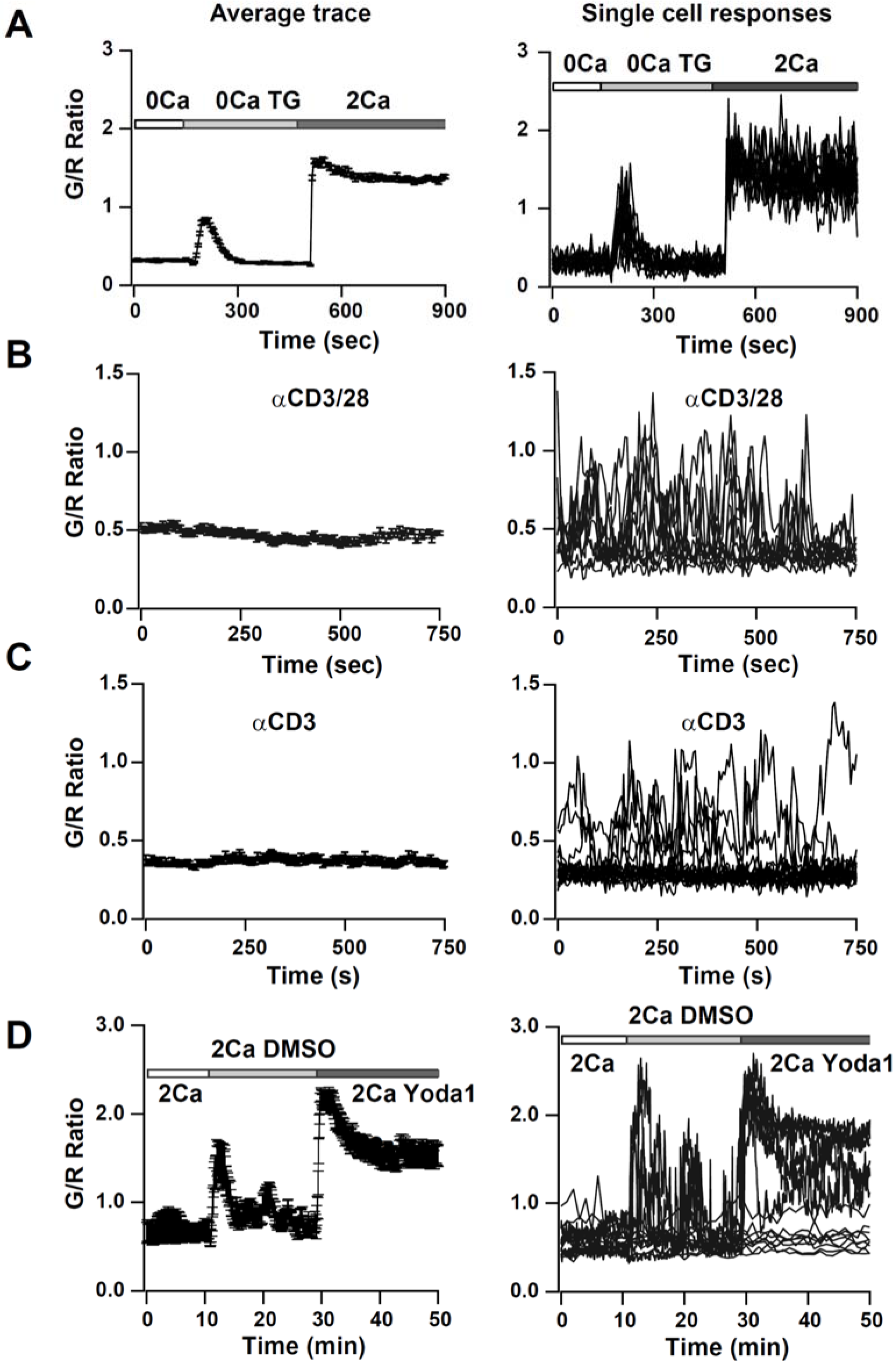
Ca^2+^ signals in activated CD4^+^ T cells from CD4-Salsa6f^+/−^ mice in response to store-depletion, TCR stimulation and Piezo1 channel activation. Average Salsa6f G/R ratios on left, representative single-cell traces superimposed on right. Experiments were done in standard Ringer solution (**A**) or in RPMI containing 2% FCS and 2 mM Ca^2+^ (**B-D**). **(A)** Store-operated Ca^2+^ entry (SOCE) in CD4^+^ T cells (n = 86 cells), induced by depleting ER Ca^2+^ stores with TG in Ca^2+^-free buffer followed by re-addition of Ringer containing 2 mM Ca^2+^. **(B,C)** Ca^2+^ responses to TCR stimulation T cells plated on coverslips coated with 1 μg/ml αCD3/CD28 (**B**) or 1 μg/ml αCD3 alone (**C**) (n = 90 cells each). **(D)** Ca^2+^ elevations during shear stress induced by solution exchange followed by the Piezo1 agonist Yoda1 (15 μM) in cells plated on αCD3/28 (n = 79 cells).

In contrast, single cell analysis of cells plated on αCD3/28 to activate TCR-induced signaling revealed a heterogeneous pattern of activation with cells showing asynchronous Ca^2+^ oscillations of varying frequencies and time-widths with a percentage of cells not responding at all (**Figure 7B**, **Video 4**). This is in contrast to a single transient Ca^2+^ peak in response to soluble αCD3/28 (data not shown). The average elevation of cytosolic Ca^2+^ was significantly above the resting levels, but below that of TG-induced SOCE. Past studies have attributed these Ca^2+^ oscillations to SOCE from repetitive opening and closing of Orai1 channels allowing Ca^2+^ to enter T cells in a periodic and asynchronous manner (Lewis and Cahalan 1989, Dolmetsch and Lewis 1994), unlike with TG treatment. Cells plated on αCD3 alone also showed rhythmic oscillatory Ca^2+^ signals; however, the percentage of responding cells was significantly lower than with αCD3/28, resulting in a lower average signal (**Figure 7C**, **Video 5**). These results suggest that co-stimulatory signaling through CD28 is essential to maintain TCR signaling, in alignment with previous observations (Chen and Flies 2013).

Finally, we focused on a novel Ca^2+^ signaling pathway as yet unreported in T cells. The Piezo family of mechanosensitive channels plays a vital role in cell motility and development (Nourse and Pathak 2017). It is not known whether these channels are expressed, and if they play any role in immune cell function. Recently, Yoda1, a selective small molecule activator of Piezo1 was identified through a drug screen (Syeda, Xu et al. 2015). We examined activation of mechanosensitive channels in cells that were plated on αCD3/28 coated glass coverslips, by flowing external solution rapidly past the cells. Perfusion of media alone produced a transient rise in the cytosolic Ca^2+^ signal (**Figure 7D**). We then tested responses to Yoda1 to assess whether Piezo1 channels can be activated in T cells and found robust and sustained Ca^2+^ signals evoked by Yoda1. Ca^2+^ responses to TG and Yoda1 responses in naïve T cells were very similar to those observed in 2-day activated T cells (**Figure 7-figure supplement 1**). In summary, for the first time, we show Ca^2+^ signals in T cells in response to activation of Piezo1 channels. These results illustrate the utility of Salsa6f for screening agents that modulate Ca^2+^ signaling in T cells and open the possibility for further exploration of the functional role of Piezo1 channels in T cell function.

### TCR-induced Ca^2+^ signaling in helper T cell subsets

We also monitored Ca^2+^ signaling in response to TCR activation with αCD3/28 in various subsets of T cells from CD4-Salsa6f^+/−^ mice, including naïve T cells, Th17 cells and iTregs **(Figure 8A-C)**. All subtypes of T cells responded to plate-bound stimulation of αCD3/28 with oscillatory changes in their cytosolic Ca^2+^ levels, very similar to the Ca^2+^ responses seen in 2 day-activated T cells shown in the previous figure. Furthermore, as with 2 day-activated T cells, responses were heterogeneous, with cells showing multiple peaks of varying width and amplitude, occasional sustained signals and a variable percentage of non-responders. While the overall average responses were not very different between the three subtypes examined here, single cell responses in Th17 cells and iTregs showed higher amplitude signals than naïve T cells, but with a greater percentage of non-responding cells. Taken together, we conclude that the CD4-Salsa6f^+/−^ mouse opens up new avenues to study the fundamental nature of Ca^2+^ signals in T cell subsets, generated in response to variety of stimuli and to explore the relationship between types of Ca^2+^ signals and specific downstream functions.

**Figure 8.**
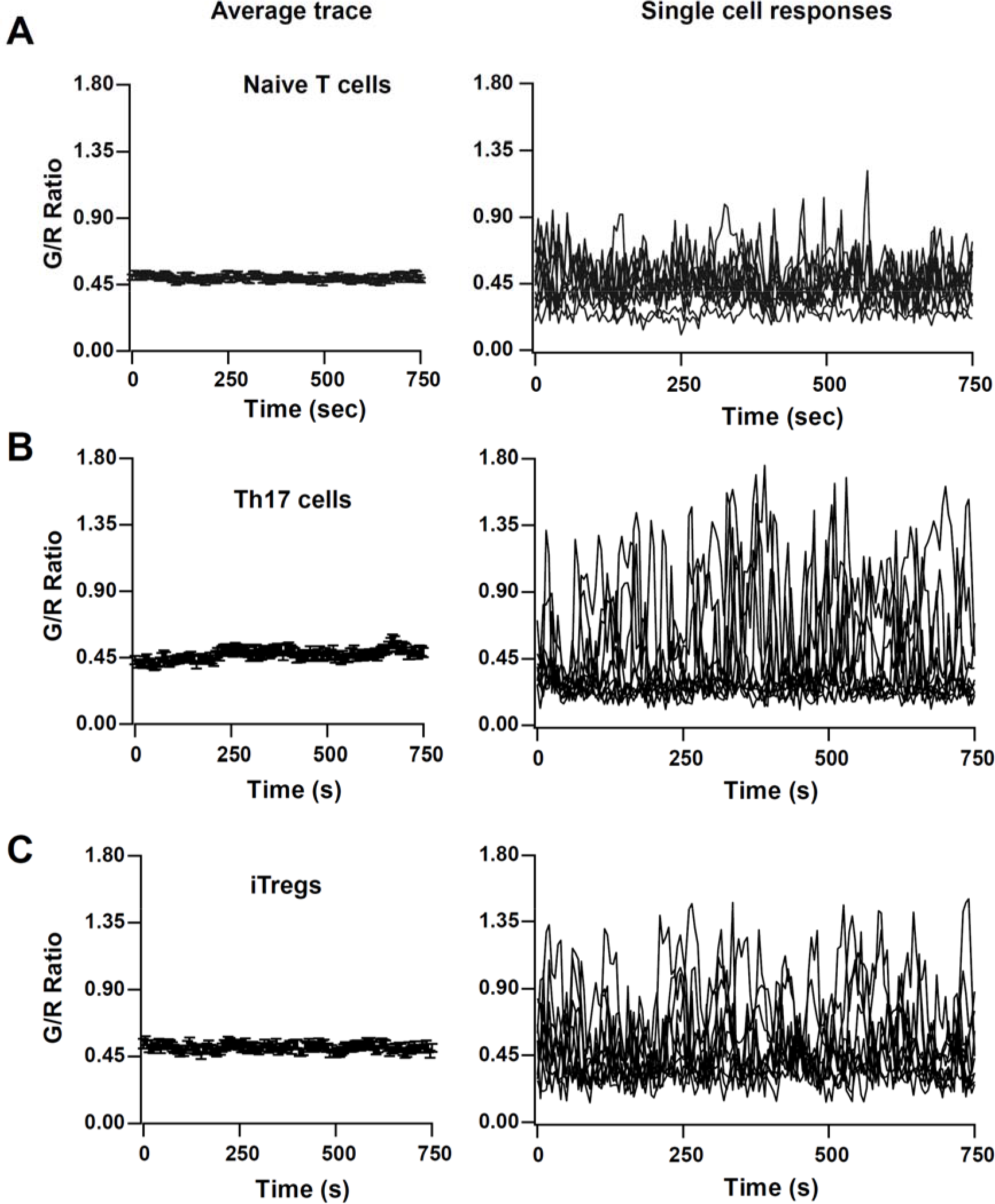
TCR induced Ca^2+^ signals in T cell subsets from CD4-Salsa6f^+/−^ mice. Average and representative single-cell Ca^2+^ traces from confocal time-lapse microscopy showing changes in Salsa6f green/red (G/R) ratio in naïve T cells (**A**), 5 day differentiated Th17 cells (**B**), and 5 day differentiated iTregs (**C**) plated on 1 μg/mL αCD3/28. (n = 90 cells from 2 - 3 experiments each).

### Two-photon microscopy of CD4-Salsa6f^+/+^ T cells in lymph node

After establishing Salsa6f as a robust reporter of cytosolic Ca^2+^, we imaged lymph nodes from homozygous CD4-Salsa6f^+/+^ mice using 2-photon microscopy under steady-state conditions. Over time, sporadic T cell-sized green fluorescent signals were seen, and the pattern of fluorescence observed is consistent with the cytosolic localization of the Salsa6f indicator, which is excluded from the nucleus (**Figure 9A-C**). Numerous small, bright, and transient green fluorescent signals about 2 μm^2^ in area were also observed (**Figure 9B,D**; **Video 6**). We termed these fluorescent transients “sparkles”, because during rapid playback of time-lapse image streams cells appear to sparkle (**Video 6**). The existence of sparkles was surprising, because sparkles are too small to result from cell-wide elevations of Ca^2+^, and T cells lack extended cellular processes, like neurons, that confine Ca^2+^ responses.

**Figure 9.**
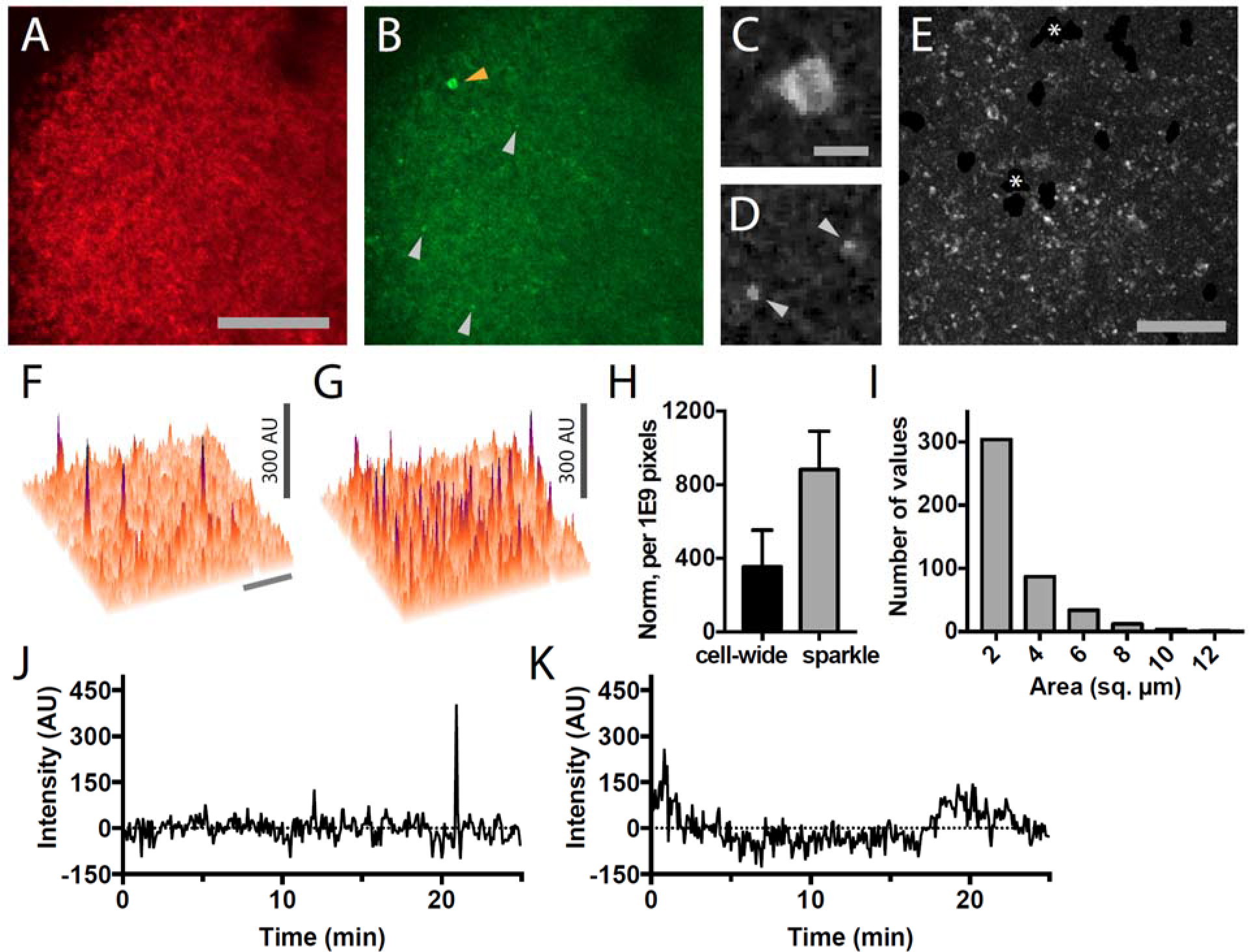
Lymph nodes from CD4-Salsa6f^+/+^ mice exhibit cell-wide and subcellular Ca^2+^ signals. (**A**) Median filtered, maximum intensity projection of a red channel image from a single time point of an explanted lymph node from a CD4-Salsa6f^+/+^ mouse. (**B**) Green channel image corresponding to **A**. Orange arrowhead indicates cell-wide Ca^2+^ signal and gray arrowheads indicate smaller, local transient Ca^2+^ signals. (**C, D**) Enlargements of cell-wide (**C**) and local (**D**; gray arrowheads) Ca^2+^ signals. Note the lower fluorescence intensity in the center of the cell in C due to exclusion of Salsa6f from the nucleus. (**E**) Maximum intensity projection of 214 green channel time points (every 11.5 seconds over 41 minutes) showing hundreds of small local Ca^2+^ signals. Green channel image series was red channel subtracted and cropped from **B**. Asterisks indicate regions containing autofluorescent cells that have been cropped out. (**F, G**) Surface plot of maximum green channel intensity over two (**F**) and 50 (**G**) consecutive time points. Note the presence of four (**F**) and dozens (**G**) of small, discrete, high intensity peaks of similar intensity. (**H**) Bar graph of relative frequencies of cell-wide and local Ca^2+^ signals. (**I**) Frequency distribution of the area of local Ca^2+^ signals. Scale bar in **A** is 100 μm (applies to **B**); scale bar in **C** is 10 μm (applies to **D**), scale bars in **E** and in **F** are 50 μm (applies to **G**). (**J**) Trace of fluorescence intensity over 25 minutes at the location of a transient subcellular Ca^2+^ signal (one time point every 5 seconds). (**K**) Trace of fluorescence intensity of a putative cell process from an autofluorescent cell drifting in the image field.

Since T cells move rapidly and are not uniformly distributed in lymph nodes, we developed an image processing approach to minimize fluctuations in background fluorescence in order to sensitively identify cell-wide Ca^2+^ signals and sparkles (**Figure 9-figure supplement 1**). A key feature of the Salsa6f fusion protein is the one-to-one correspondence of tdTomato and GCaMP6f, and we used this to estimate and subtract out fluctuations in green background fluorescence due to cell movement and distribution. After processing, sparkles were found to occur widely across the imaging field and many had similar intensities (**Figure 9E-G**). The brightness of sparkles and the uniformity of background fluorescence allowed us to use a stringent 5.4 SD threshold to systematically identify bright sparkles; hundreds (mean of 6.5 SD above background) were observed in each 25-minute imaging session (one image stack every five seconds), whereas less than one was expected to occur by chance. Sparkles were also more frequent than cell-wide transients (**Figure 9H**). Most sparkles were between the defined minimum 1.4 μm^2^ and 3 μm^2^ in area (median of 1.9 μm^2^, 95% CI of 1.9-2.3 μm^2^; n=441 sparkles from 3 cells; **Figure 9I**). Sparkles were typically found in one or two consecutive frames (**Figure 9J**). Sparkle trace shape differs from that expected for autofluorescent cell processes drifting into the imaging field (**Figure 9K**). Taken together, these observations suggest that sparkles correspond to local Ca^2+^ signals restricted to small subcellular domains of T cells migrating through the lymph node.

To establish that T cells labeled in CD4-Salsa6f^+/+^ mice exhibit subcellular Ca^2+^ signals, we adoptively transferred CD4-Salsa6f^+/+^ T cells into wild type recipients so these cells could be viewed in isolation. Sparkles were identified in time lapse images of lymph nodes, and when traced back they were found to originate in red fluorescent cells in most cases. Thus, sparkles correspond to restricted subcellular domains of elevated Ca^2+^ in Salsa6f CD4^+^ T cells. Cell movement was used to define the front and back of labeled T cells for mapping the subcellular location of Ca^2+^ signals. Local Ca^2+^ signals were most frequently found in the back of motile T cells (**Figure 10 A-E**). Green-red channel ratiometric images, enabled by Salsa6f labeling, confirmed cytosolic localization patterns (**Figure 10D**). Local Ca^2+^ signals were found less frequently in the front, front and back, and sides of motile T cells (**Figure 10 F-H**). In a companion paper, we use Salsa6f transgenic mice to consider the relationship between Ca^2+^ signals, both cell-wide and local, and T cell motility in the lymph node.

**Figure 10.**
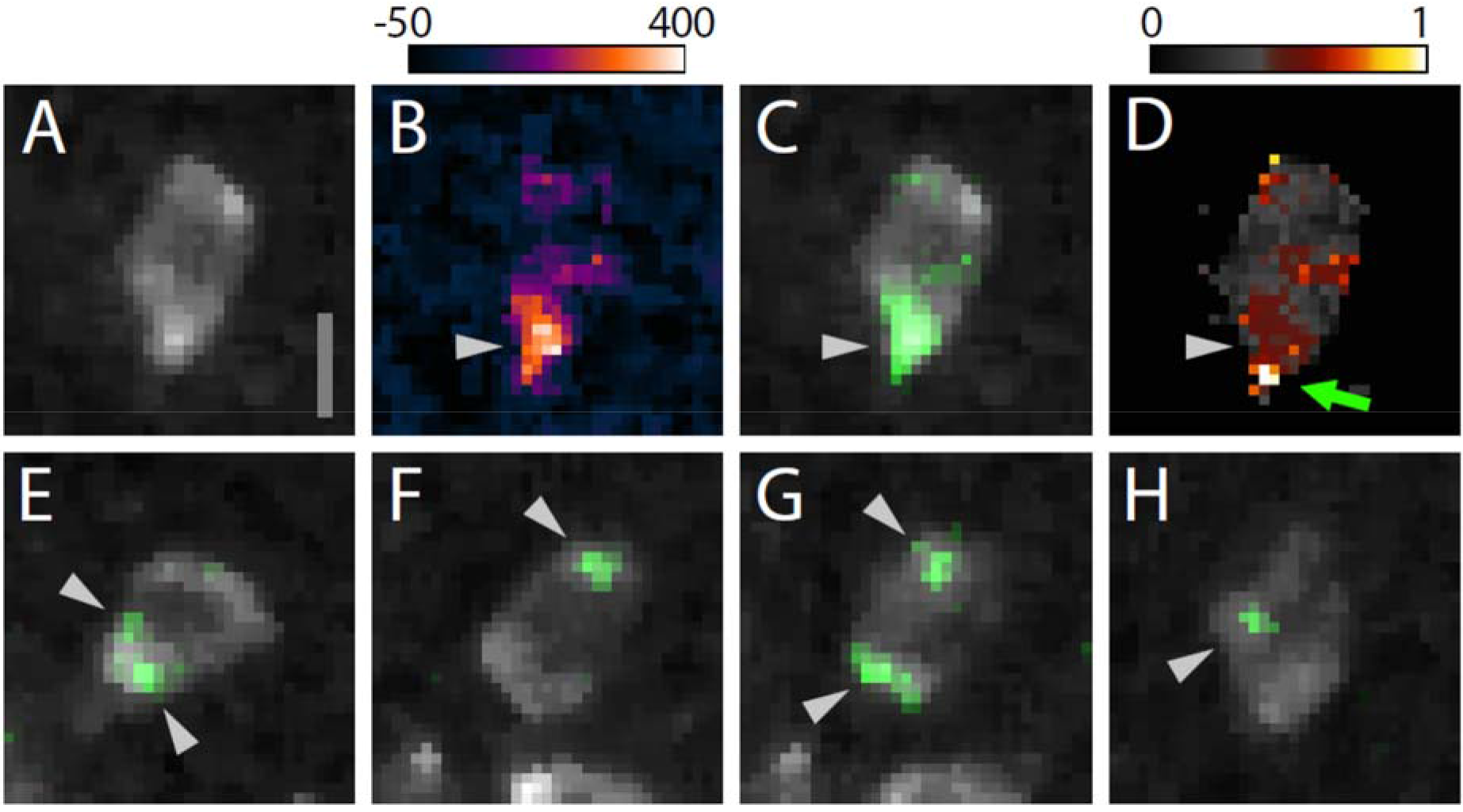
Subcellular Ca^2+^ signals map to different regions of motile CD4-Salsa6f^+/+^ T cells. (**A-D**) CD4-Salsa6f^+/+^ T cell imaged in a wild type lymph node after adoptive transfer. (**A**) Red channel fluorescence image. (**B**) Corresponding pseudocolored green channel image. (**C**) Corresponding composite image of gray pseudocolored red channel image with green channel image. (**D**) Ratiometric image of the green divided by the red channel fluorescence image. Gray arrowheads denote a local Ca^2+^ signal at the back of motile T cell and the green arrow denotes a point of relatively high Ca^2+^ concentration at the extreme back of the cell. Look-up table for **B** corresponds to Arbitrary Units; look-up table for **D** corresponds to green-to-red ratio. (**E-H**) Adoptively transferred CD4-Salsa6f^+/+^ T cells displaying local Ca^2+^ signals at different subcellular locations. (**E**) Two local Ca^2+^ signals at the back of the cell. (**F**) Local Ca^2+^ signal at the front of a different cell. (**G**) Same cell as in F five seconds later displaying local Ca^2+^ signals at both the front and back. (**H**) Different cell displaying local Ca^2+^ signal at the side. Location of local Ca^2+^ signals indicated by gray arrowheads. For all, cells are oriented with their front toward the top of the image. Scale bar in A is 5 μm (applies to **B-H**).

## Discussion

We introduce Salsa6f, a novel, ratiometric genetically-encoded Ca^2+^ probe. Salsa6f is a fusion of the high performing green fluorescent GECI GCaMP6f and the bright red fluorescent tdTomato. This simple modification imparts powerful capabilities, which include tracking cells in the absence of Ca^2+^ signaling, ratiometric imaging to eliminate motility artifacts, and convenient single-wavelength femtosecond excitation for two-photon microscopy. Salsa6f addresses a key weakness of single fluorescent protein-based GECIs by enabling tracking of motile cells and identification of cell morphology, even at basal Ca^2+^ levels when GCaMP6f fluorescence is very weak. We further generated a transgenic reporter mouse with Cre-dependent expression of Salsa6f, enabling Ca^2+^ signals to be imaged in specific, genetically defined cell types. Transgenic expression of Salsa6f brings the power of ratiometric chemical Ca^2+^ indicators to imaging cellular Ca^2+^ signals amid the complex tissue environments found in vivo.

Salsa6f preserves the exceptional performance of GCaMP6f, which in the presence of high levels of Ca^2+^ is as bright as the standard high performing green fluorescent protein, EGFP (Chen, Wardill et al. 2013). We find that Salsa6f possesses a dynamic range similar to GCaMP6f as well, and both are superior to FRET-based GECIs (Heim, Garaschuk et al. 2007, Thestrup, Litzlbauer et al. 2014). Salsa6f’s Ca^2+^ affinity, 160-300 nM, is well suited to detecting a variety of cellular Ca^2+^ signals. Inclusion of tdTomato in Salsa6f enables ratiometric imaging, calibration, and measurement of Ca^2+^ concentrations within cells. Salsa6f is uniformly distributed throughout the cytosol; its exclusion from the nucleus provides reliable and selective reporting of cytosolic Ca^2+^ signaling. This is in contrast to the recently developed CD4-Cre 5GtdT^+/−^ mouse strain in which the tdTomato is found throughout the cell but the separately expressed GCaMP5G is excluded from the nucleus (Gee, Smith et al. 2014). Finally, Salsa6f expression is non-perturbing; we saw no effects of Salsa6f expression in CD4^+^ immune cells with respect to cellular phenotype, cell proliferation, differentiation, and, in our companion paper, homing and T cell motility.

We have created a transgenic mouse strain in which Salsa6f is expressed under genetic control using the Rosa26-Cre recombinase system, and we have used this system to label immune cells that express CD4. Salsa6f labeling enables readout of cytosolic Ca^2+^ dynamics in T cells in vitro with high dynamic range without the handling and potential toxicity associated with loading of chemical Ca^2+^ indicators. Salsa6f was used to detect Ca^2+^ influx due to direct activation of SOCE, TCR stimulation, and Piezo1 channel opening, detected in T cells for the first time to our knowledge. We also detected differences in patterns of Ca^2+^ signaling between naïve T cells, Th17 cells, and iTregs. These experiments demonstrate the sensitivity, brightness, uniformity of labeling, and ease of detecting dynamic Ca^2+^ signals using Salsa6f.

A primary advance of this work is to take the in vitro capabilities of an excellent Ca^2+^ indicator and bring these into the realm of in vivo imaging. Within tissues, cells at a given position exhibit differences, ranging from subtle to dramatic, in morphology, connectivity, and molecular profile. The red fluorescence of Salsa6f, combined with genetic Salsa6f labeling, associates these characteristics with readout of cellular Ca^2+^ signaling. Red fluorescence of Salsa6f is well excited by the same wavelength used for 2-photon imaging of GCaMP6f, 920 nm, enabling imaging hundreds of micrometers deep into lymph nodes. The immune system poses additional challenges for imaging because the constituent cells are highly motile in lymphoid organs. Indeed, direct cell interactions of motile immune cells form the basis of immune surveillance. We were able to identify red fluorescent Salsa6f T cells easily in intact lymph nodes upon adoptive transfer. Our images reveal uniform red fluorescence labeling by Salsa6f with clear subcellular morphology in imaging sessions encompassing hundreds of time lapse images. Images of lymph nodes from CD4-Salsa6f^+/+^ mice display dozens to hundreds of cell-wide Ca^2+^ responses even in the absence of antigen. These capabilities indicate that Salsa6f transgenic mice could be used to associate Ca^2+^ signaling with T cell behavior in vivo.

Salsa6f offers the opportunity not only to record fluctuations in relative Ca^2+^ levels over time, but also to read out Ca^2+^ concentrations within cells. We have measured the affinity of Salsa6f in intact cells; use of this approach will allow other microscope systems to be calibrated for measuring absolute Ca^2+^ concentrations with Salsa6f. Knowledge of absolute Ca^2+^ concentrations is necessary to develop quantitative models of Ca^2+^ signaling and cell behavior. Indeed, we demonstrate that clear Salsa6f ratio images can be generated from motile T cells in intact lymph nodes.

Our Salsa6f transgenic mouse line enables more sophisticated experimental approaches. One is the ability to detect rare Ca^2+^ signaling events. The high brightness and dynamic range of modern GECIs like Salsa6f contribute to detection of rare Ca^2+^ signaling events inside intact tissues or even whole transgenic animals (Kubo, Hablitzel et al. 2014, Portugues, Feierstein et al. 2014). Detecting rare events is made harder by inhomogeneities in cell populations of the lymph node as well as the movement of immune cells therein. Because of the one-to-one correspondence of tdTomato and GCaMP6f in Salsa6f, we were able to estimate and subtract resting GCaMP6f fluorescence even in motile cells. This approach substantially improves the uniformity of the fluorescence background upon which rare Ca^2+^ signaling events are detected. Reliable and uniform cytosolic labeling contributes as well. Combined, these factors enabled us to detect not only sporadic cell-wide Ca^2+^ elevations, but also unexpectedly sparkles, much smaller sporadic local Ca^2+^ signals. The sensitivity and resolution of these images are sufficient to map local signals from intact lymph nodes to sub-regions of T cells, which are some of the smallest cells of the body. Moreover, while we focused upon the brightest local Ca^2+^ signals to demonstrate their existence, we expect that Salsa6f will enable lower intensity Ca^2+^ signals to be linked to subcellular mechanisms and, ultimately, resulting cell behaviors. In a companion paper we relate Ca^2+^ signals detected by Salsa6f, both global and local, to T cell motility in the lymph node.

In conclusion, here we demonstrate the utility our Ca^2+^ indicator Salsa6f, and the transgenic mouse line which expresses Salsa6f, for studies of immune cell function. Use of Salsa6f improves assays of Ca^2+^ signaling in immune cell function, both in purified cell populations as well as in vivo, and for the first time to our knowledge, we detected Ca^2+^ influx associated with Piezo1 channel opening in T cells. Ca^2+^ signals were detected in T cells in lymph nodes under basal steady state conditions in the absence of antigen. Many of these Ca^2+^ signals were localized to sub-regions of T cells. Finally, we anticipate that this new probe of Ca^2+^ signaling will be widely applicable for studies of other cell types in other tissues.

## Acknowledgments

We acknowledge the UC Irvine Transgenic Mouse Facility for support in making the transgenic mouse, Dr. Grant MacGregor from the UC Irvine Department of Developmental and Cell Biology for excellent advice, and Dr. Jennifer Atwood of the Flow Core Facility supported by the UC Irvine Institute of Immunology. We also thank Andy Yeromin for advice on curve-fitting. This work was supported by an R21 grant AI117555, and two RO1 grants, NS14609 and AI121945, from the National Institutes of Health to MDC, and by a postdoctoral fellowship from the George E. Hewitt Foundation for Medical Research (A.J.).

## Methods

### GECI screening and Salsa6f plasmid generation

Plasmids encoding GECIs (GECO and GCaMP6) were obtained from Addgene for screening in live cells. Each probe was cotransfected with Orai1 and STIM1 into HEK 293A cells using Lipofectamine 2000 (Invitrogen, Carlsbad, CA) for 48 hr before screening on an epifluorescence microscope. For construction of Salsa6f, a plasmid for tdTomato (Addgene, Cambridge, MA) and the pEGP-N1 vector (Clontech, Mountain View, CA) was used as a backbone. GCaMP6f was amplified via PCR with N- and C-terminal primers (5’ CACAACCGGTCGCCACCATGGTCGACTCATCACGTC 3’ and 5’ AGTCGCGGCCGCTTTAAAGCTTCGCTGTCATCATTTGTAC 3’) and ligated into pEGFP-N1 at the AgeI/NotI sites to replace the eGFP gene, while tdTomato was amplified via PCR with N- and C-terminal primers (5’ ATCCGCTAGCGCTACCGGTCGCC 3’ and 5’ TAACGAGATCTGCTTGTACAGCTCGTCCATGCC 3’) and ligated into the backbone at the NheI/BglII sites. An oligo containing the V5 epitope tag was synthesized with sense and antisense strands (5’ GATCTCGGGTAAGCCTATCCCTAACCCTCTCCTCGGTCTCGATTCTACG 3’ and 5’ GATCCGTAGAATCGAGACCGAGGAGAGGGTTAGGGATAGGCTTACCCGA 3’) and ligated into the backbone at the BglII/BamHI sites, linking tdTomato to GCaMP6f and creating Salsa6f. The amplified regions of the construct were verified by sequencing (Eton Bioscience Inc., San Diego, CA). This plasmid, driven by the CMV promoter, was used for transient transfections in HEK 293A cells with Lipofectamine 2000 and in primary human T cells with Amaxa Nucleofection.

### Transgenic mouse generation and breeding

The transgenic cassette in **Figure 2B** was generated by inserting Salsa6f, from the plasmid described above, into the Ai38 vector (Addgene Plasmid #34883) and replacing GCaMP3. The final targeting vector included the CAG (cytomegalovirus early enhancer/chicken β-actin) promoter, an LSL sequence with LoxP-STOP-LoxP, the Salsa6f probe (tdTomato-V5-GCaMP6f), the woodchuck hepatitis virus posttranscriptional regulatory element (WPRE), and a neomycin resistance gene (NeoR), all flanked by 5’ and 3’ Rosa26 homology arms of 1.1 and 4.3 kb. The targeting vector was linearized with PvuI and electroporated into JM8.N4 mouse embryonic stem (ES) cells of C57BL/6N background. Following selection with G418, clones were screened by Southern blotting after digestion with HindIII for the 5’ end or BglI for the 3’ end. Four correctly targeted clones were expanded and checked by chromosome counting, then two clones with >90% euploidy were further expanded and injected into C57BL/6J blastocysts for implantation into pseudopregnant foster mothers. Presence of the Salsa6f transgenic cassette was detected in the resulting chimeric pups by PCR screening for the *Nnt* gene, as the initial JM8.N4 ES cells are *Nnt*^+/+^ while the C57BL/6J blastocysts are *Nnt*^−/−^. Finally, positive chimeras were bred to R26ΦC31o mice (JAX #007743) to remove the neomycin resistance gene flanked by AttB and AttP sites in the original transgenic cassette, and to produce Salsa6f^LSL/−^ F1 founders carrying the allele for LSL-Salsa6f at the Rosa26 locus. These F1 founders were then bred to homozygosity to generate Salsa6f^LSL/LSL^ mice, and subsequently crossed to homozygotic CD4-Cre mice (JAX #017336) to generate CD4-Salsa6f^+/−^ mice expressing Salsa6f only in T cells. CD4-Salsa6f^+/−^ mice were further bred to generate homozygotic CD4-Salsa6f^+/+^ mice for increased Salsa6f expression and fluorescence.

### T cell proliferation and differentiation

*For T cell proliferation*: CD4 T cells were isolated from spleen and lymph nodes of 6-10 week old mice using negative selection (StemCell Technologies, Cambridge, MA). CellTrace Violet (CTV)-labeled T cells were co-cultured with αCD3/CD28 coated dynabeads (Life Technologies Corp., Grand Island, NY) at 1:1 ratio according to the manufacturer’s protocol in a U bottom 96 well plate. *For T cell differentiation*: Naïve CD4 T cells were differentiated on activating polystyrene surface (Corning Inc., Corning, NY) with plate-bound αCD3 (2.5 μg/ml) and αCD28 (2.5 μg/ml) in the presence of cytokines for 6 days (Yosef, Shalek et al. 2013). For Th1 differentiation: 25 ng/mL rmIL-12 (BioLegend, San Diego, CA), 10 μg/mL αmouse IL4 (Biolegend). For Th17 differentiation: 2.5 ng/mL rhTGF-β1 (Tonbo Biosciences, San Diego, CA), 50 ng/mL rmIL-6 (Tonbo Biosciences), 25 ng/ml rmIL-23 (Biolegend), and 25 ng/ml rmIL-β1 (Biolegend). For iTreg differentiation: 10 ng/mL rhTGF-β1, 100 units/mL of rmIL-2 (Biolegend), 5 μM Retinoic Acid (Sigma, St. Louis, MO).

### Flow cytometry

CTV dilution assay was performed in live cells (Fixable Viability Dye eFluor^®^ 780 negative gating; Thermofisher Scientific Inc., Grand Island, NY). To detect intracellular cytokines, 6 day differentiated cells were stimulated in with 25 ng of phorbol 12-myristate 13-acetate (PMA), 1 μg ionomycin (Sigma), and monensin (Golgistop^®^ BD biosciences) for 4 hr at 37 °C. Dead cells were labeled with Ghost dye 780 (BioLegend), then washed, fixed, permeabilized using FoxP3 staining buffer set (Thermofisher Inc).

The following antibodies were used to detect intracellular cytokines: IL-17A-APC (clone TC11-18H10.1, BioLegend); IFNγ-Pacific Blue (clone XMG1.2, BioLegend); Foxp3-PE (clone FJK16s, Thermofisher Scientific Inc.); in permeabilization buffer (eBioscience). Data were acquired using NovoCyte flow cytometer (ACEA Biosciences) and analyzed using FlowJo.

### T-cell preparation for live cell imaging

CD4 T cells were activated by plating on 6 well plates coated overnight with 2.5 μg/mL αCD3/αCD28 (Invivogen, San Diego, CA) at 4° C. Cells were cultured in RPMI medium (Lonza) containing 10% FCS, L-glutamine, Non-essential amino acids, Sodium pyruvate, β-mercaptoethanol and 50 U/mL of IL-2 at 37° C in 5% CO_2_ incubator. Following 2 days of culture, cells were plated on either poly-L-lysine or 1 μg/mL α-CD3/α28 coated 35mm glass chambers (Lab-Tek, Thermofisher Inc.) for imaging. RPMI medium with 2% FCS and L-glutamine containing 2 mM Ca^2+^ was used for imaging experiments. For experiments involving calibration and characterization of the Salsa6f probe in CD4-Salsa6f^+/−^ cells, Ringer solution containing various concentrations of Ca^2+^ was used. For Ca^2+^ imaging of different T cell subsets, Th17 cells and iTregs were differentiated as described above.

### Confocal imaging and analysis

For Ca^2+^ imaging of CD4^+^ T cells from CD4-Salsa6f mice, we used an Olympus Fluoview FV3000RS confocal laser scanning microscope, equipped with high speed resonance scanner and the IX3-ZDC2 Z-drift compensator (Olympus Corp., Waltham, MA). Diode lasers (488 and 561 nm) were used for excitation, and two high-sensitivity cooled GaAsP PMTs were used for detection. Cells were imaged using the Olympus 40x silicone oil objective (NA 1.25), by taking 5 slice z-stacks at 2 μm/step, at 5 sec intervals, for up to 20 min. Temperature, humidity, and CO_2_ were maintained using a Tokai-Hit WSKM-F1 stagetop incubator. Data were processed and analyzed using Imaris and ImageJ software. Calcium imaging experiments were done at 37° C on 2 day-activated CD4^+^ T cells from CD4-Salsa6f^+/−^ mice, unless otherwise indicated. Salsa6f calibration experiments were done at room temperature.

### Two-photon microscopy

Lymph nodes images were acquired using a custom-built two photon microscope based on Olympus BX51 upright frame, Motorized ZDeck stage (Prior, Rockland, MA), with excitation generated by a tunable Chameleon femtosecond laser (Coherent, Santa Clara, CA) (Miller, Wei et al. 2002). The following wavelengths were used to excite single or combination of fluorophores: 920 nm to excite tdTomato and GCaMP6f; 1040 nm to excite tdTomato alone. 495 nm and 538 nm dichroic filters were arranged in series to separate blue, green and red signals. Two-photon excitation maxima of tdTomato and GCaMP6f are 1040 and 920 nm, respectively (Drobizhev, Makarov et al. 2011, Chen, Wardill et al. 2013). Using 1040 nm excitation, tdTomato signals were readily detected up to 300 μm depth; however, 1040 is not ideal to image Salsa6f because: 1) Collagen fibers generate second harmonic at 520 nm when excited with 1040 nm, which interferes with simultaneous detection of GCaMP6f (emission maxima, 509 nm); and 2) 1040 nm does not excite GCaMP6f (**Figure 9-figure supplement 1A, top row**). Alternatively, 920 nm optimally excites GCaMP6f, and excites tdTomato sufficiently, and Salsa6f signals were detected up to 300 μm depth, while second harmonic collagen signals (460 nm) can be easily separated into blue channel (**Figure 9-figure supplement 1A, bottom row**). Additionally, autofluorescent structures (LN resident DCs and fibroblastic reticular cells) show up as yellow bodies when excited with 920 nm, which serve as a guide to locate the T cell zone (**Figure 9-figure supplement 1B**). Therefore, 920 nm is the ideal two-photon excitation wavelength for simultaneous imaging of tdTomato and GCaMP6f as component parts of Salsa6f.

Lymph nodes were oriented with the hilum away from the water dipping microscope objective (Nikon 25x, NA 1.05). The node was maintained at 36-37°C by perfusion with medium (RPMI) bubbled with medical grade carbogen (95% O_2_ and 5% CO_2_) using a peristaltic heated perfusion system (Warner Instruments), with thermocouple-based temperature sensors placed next to the tissue in a custom built chamber. 3D image stacks of x=250 μm, y=250 μm, and z=20 or 52 μm (4 μm step size) were sequentially acquired at 5 or 11 second intervals respectively, using image acquisition software Slidebook (Intelligent Imaging Innovations) as described previously (Matheu, Othy et al. 2015). This volume collection was repeated for up to 40 min to create a 4D data set.

### Data analysis and statistical testing

Graphpad Prism was used for statistical analysis and generating figures. p values are indicated in figures: ns p> 0.05, * p < 0.05; ** p < 0.01; *** p < 0.001; and **** p < 0.0001.

### Detection of Ca^2+^ signals in lymph nodes

Stacks of 6 optical sections 4 μm apart from the T-zone of CD4-Salsa6f^+/−^ lymph nodes were acquired once every 5 sec at a resolution of 0.488 or 0.684 μm per pixel. Maximum intensity projections of 1 pixel radius median-filtered images were used for subsequent processing and analysis. Autofluorescent cells were identified by averaging the red or green time lapse image stacks and automated local thresholding (Bernsen 5 pixel radius) using the public domain image processing program ImageJ. Autofluorescent cell masks were dilated by 4 pixels, regions exhibiting less contrast and detail due to light scattering manually masked to produce the final time lapse image mask. Red (tdTomato) channel fluorescence from Salsa6f corresponding to green (GCaMP6f) channel resting state fluorescence was determined to be 5-fold higher using our standard 2-photon microscope acquisition settings. Final green images were produced by subtracting a 0.2x scaled red channel image, and subsequently subtracting out the average of all green channel time lapse images. The standard deviation (SD) of each masked green channel time lapse image stack was used to determine thresholds for local (sparkle) and cell-wide Ca^2+^ events. Thresholds for detection of local and cell-wide Ca^2+^ events were 5.4 and 2.1 SD and 1.4 μm^2^ and 25 μm^2^, respectively. Frequency of background events was calculated using a standard normal distribution with a Z-score corresponding to the average intensity of local events (6.5 SD), which was 1 in 2×10^10^ pixels (WolframAlpha).

---

**Figure 6-supplement 1.**
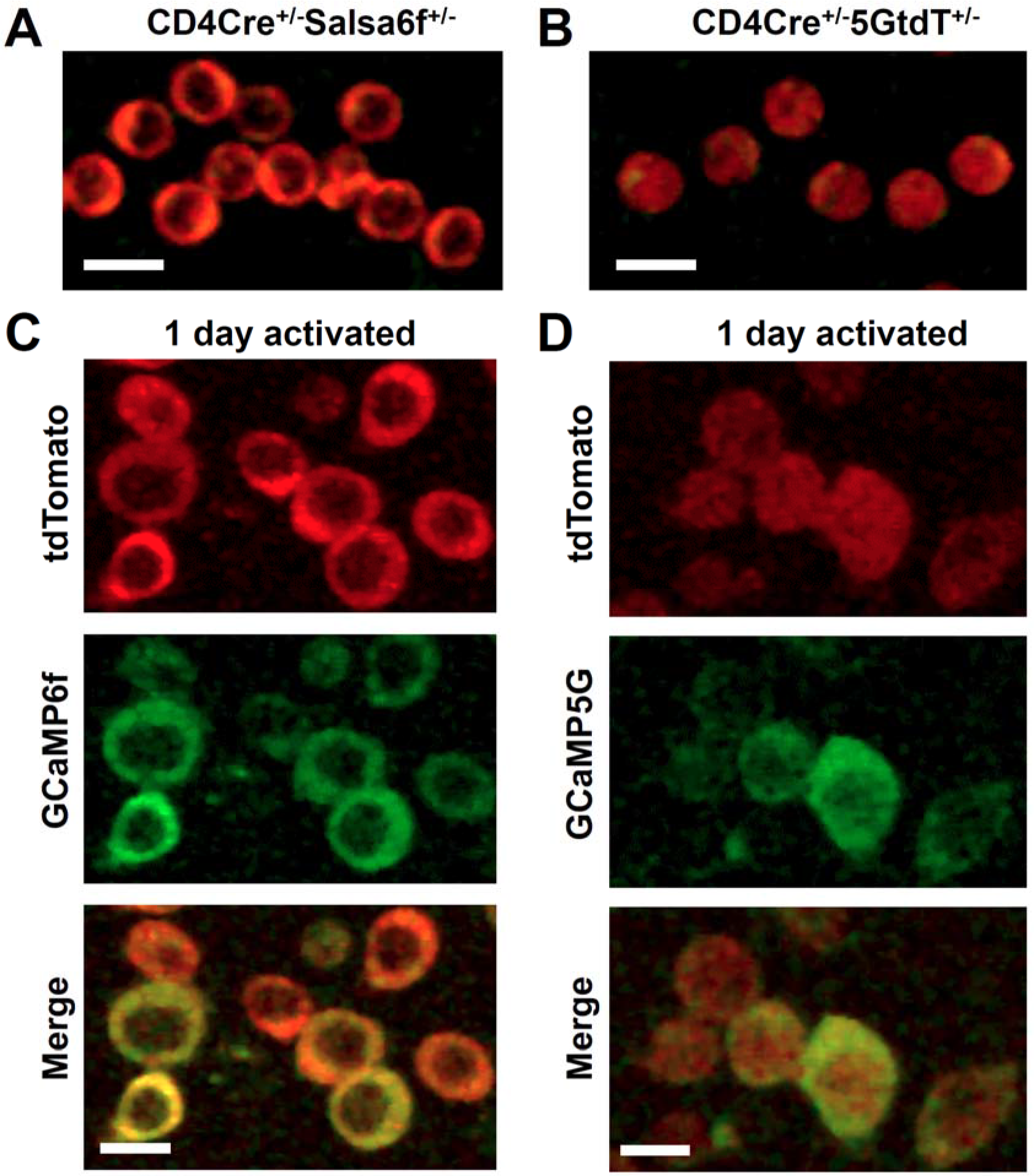
Comparison of GECI localization in CD4 T cells from Salsa6f mouse and PC::G5-tdT mouse. (**A,B**) Confocal images of CD4^+^ T cells purified from a CD4-Salsa6f^+/−^ mouse (**A**) or a CD4-Cre 5GtdT^+/-^ mouse (**B**), showing merged red and green; cells imaged at the same laser and PMT settings; scale bar = 10 μm. (**C,D**) CD4^+^ T cells purified from a CD4-Salsa6f^+/−^ mouse (**C**) or a 5G-tdT^+/−^CD4-Cre^+/−^ mouse (**D**), then activated for 24 hr on plate-bound αCD3/28 antibodies, and imaged with confocal microscopy, showing red (tdTomato), green (GCaMP6f or GCaMP5G), and merged channels; scale bar = 10 μm.

**Figure 7-figure supplement 1.**
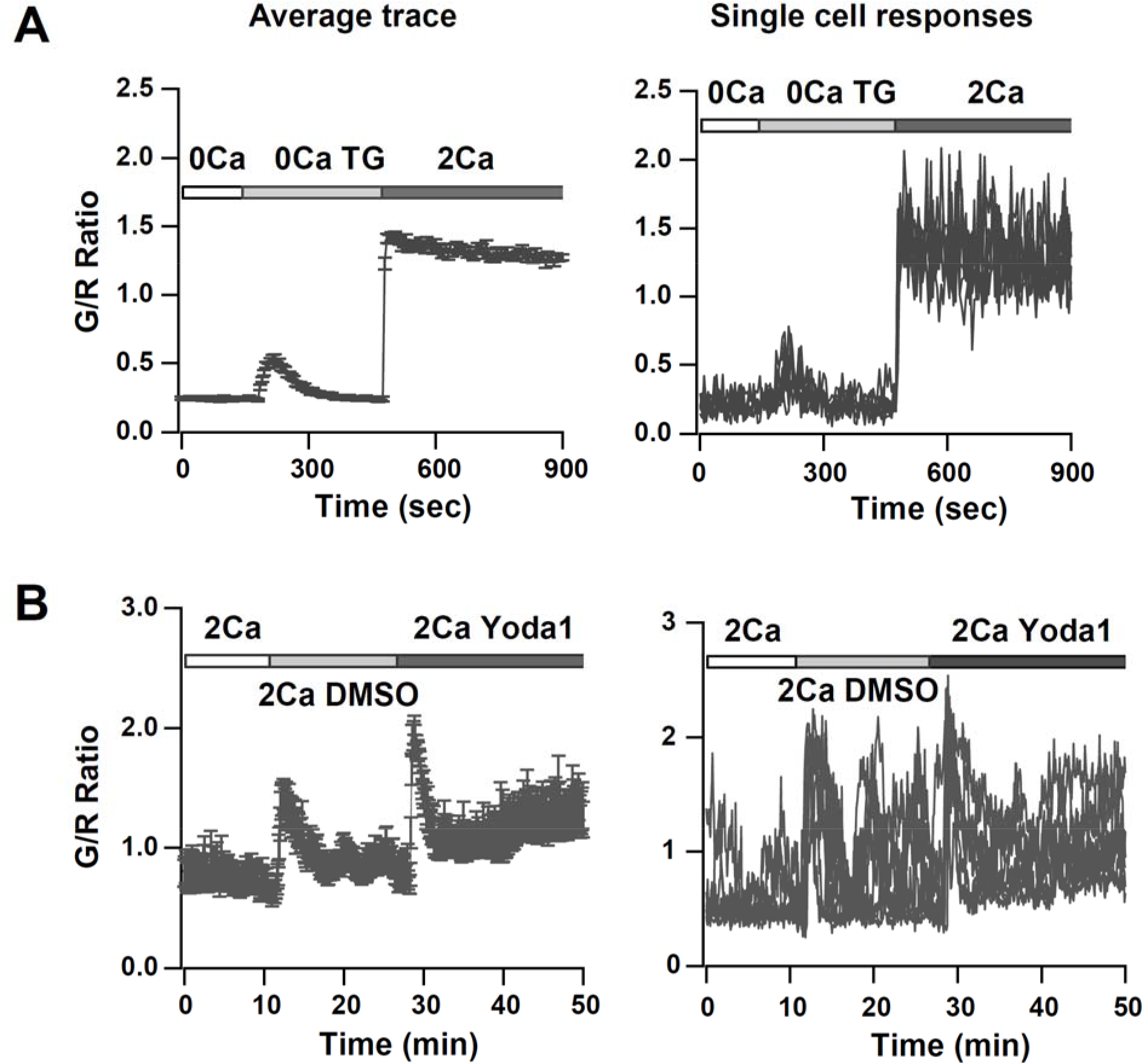
Store-operated Ca^2+^ entry and Piezo1 activation in naïve T cells from CD4-Salsa6f^+/−^ mouse. (**A**) Average (left panel) and representative single cell responses (right panel) to TG-induced SOCE (n = 96 cells). (**B**) Average (left panel) and single cell responses (right panel) to solution exchange of media alone followed by 15 μM Yoda1 (n = 53 cells).

**Figure 9-figure supplement 1.**
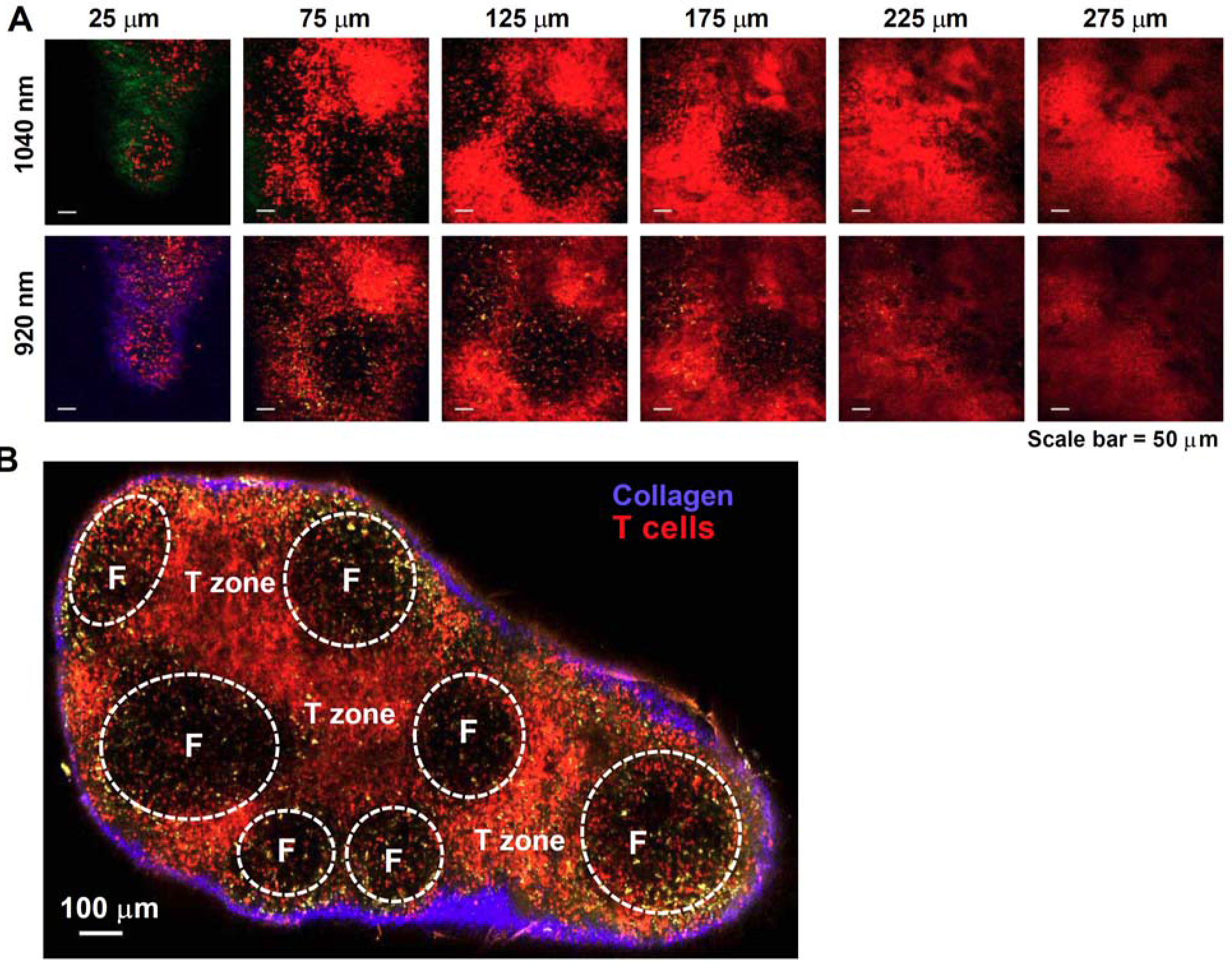
Imaging lymph nodes of CD4-Salsa6f^+/+^ homozygous mice. Cre-mediated expression of Salsa6f in CD4 T cells reveals endogenous T cell labeling in lymph node. (**A**)Two-photon images of explanted lymph node from CD4-Salsa6f^+/+^ mouse at various depths (indicated above the image); 1040 nm excitation (top, row) or 920 nm excitation (bottom row). Second harmonic signal from collagen fibers is collected in green for 1040 nm excitation and in blue for 920 nm excitation. Salsa6f cells are readily detected up to 275 μm deep. (**B**) Montage image of a CD4-Salsa6f^+/+^ lymph node at 100 μm depth, imaged using 920 nm excitation showing Salsa6f^+^ cells in red, autofluorescent structures in yellow, and the capsular boundary shown in blue (second-harmonic signal); scale bar = 100 μm.

**Figure 9-figure supplement 2.**
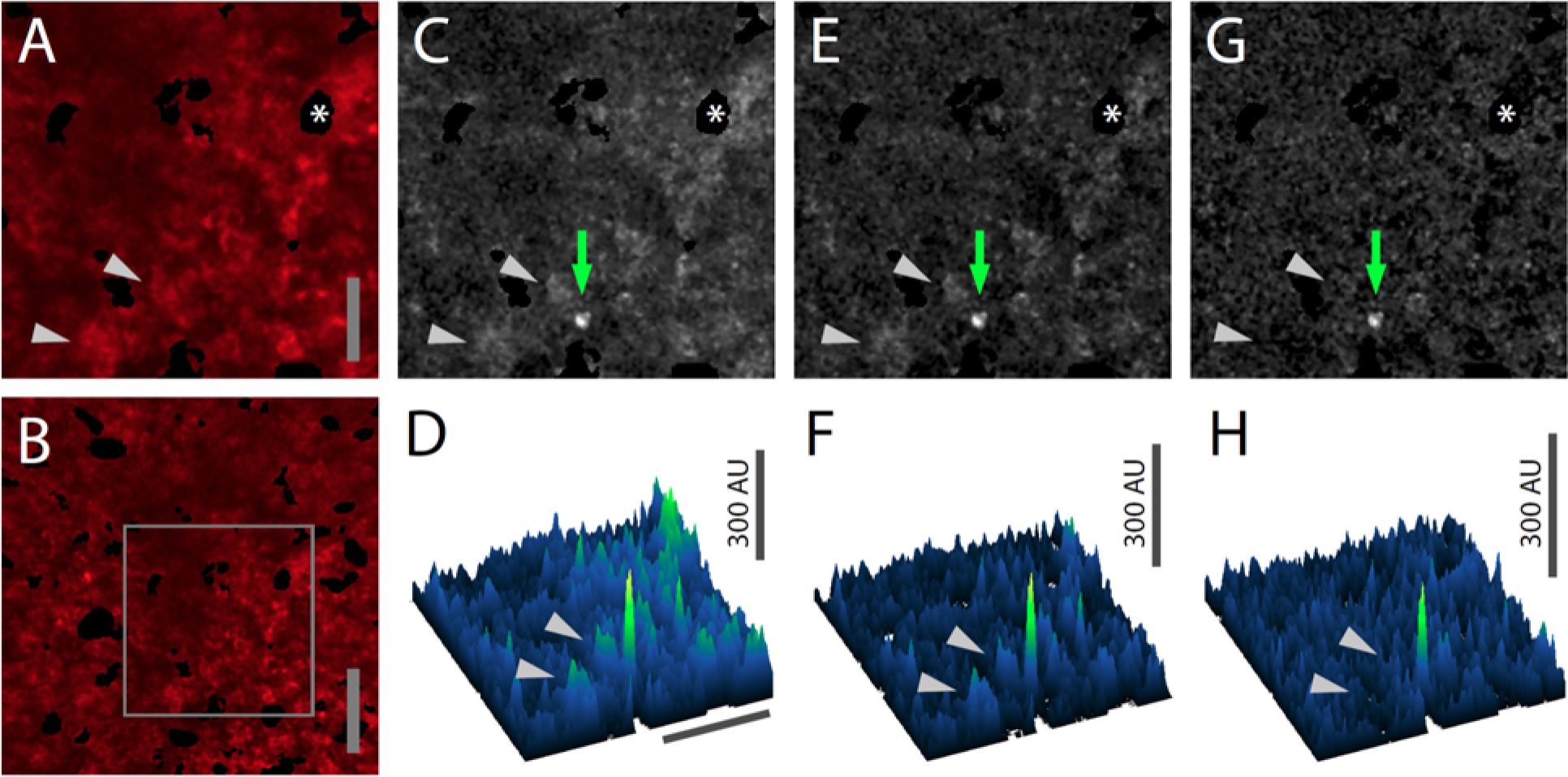
Subtraction of red channel fluorescence improves detection of Salsa6f Ca^2+^ signals. (**A, B**) Median filtered, maximum intensity projection of a red channel image from a single time point of an explanted lymph node from a CD4-Salsa6f^+/+^ mouse. Panel **A** is enlarged and cropped from panel **B** (gray rectangle). (**C, E, G**) Green channel images corresponding to **A** with different image processing protocols. (**C**) Maximum intensity projection without further processing. (**E**) Maximum intensity projection after subtraction of the average of all green channel frames. (**G**) Maximum intensity projection after subtraction of the corresponding, scaled red channel image and subtraction of the subsequent average from all green channel frames. Green arrows in **C,E,G** indicate a subcellular Ca^2+^ signal. (**D, F, H**) Surface plots corresponding to the images in **C,E,G** respectively showing the subcellular Ca^2+^ signal as a green peak. Gray arrowheads indicate nearby background cell fluorescence that is progressively removed by image processing. Asterisk indicates a region containing an autofluorescent cell that has been cropped out. Scale bar in **A** is 25 μm (applies to **C,E,G**); scale bar in **B** and the horizontal scale bar in **D** are 50 μm (applies to **F,H**).

## Proposed Cover Illustration

Montage image of a CD4-Salsa6f lymph node at 100 μm depth, imaged using 920 nm excitation showing Salsa6f^+^ cells in red, autofluorescent structures in yellow, and the capsular boundary shown in blue (second-harmonic signal); scale bar = 100 μm. CD4-Salsa6f shows abundant labeling in the T zone and interfollicular region, and scant labeling in B-cell follicles.

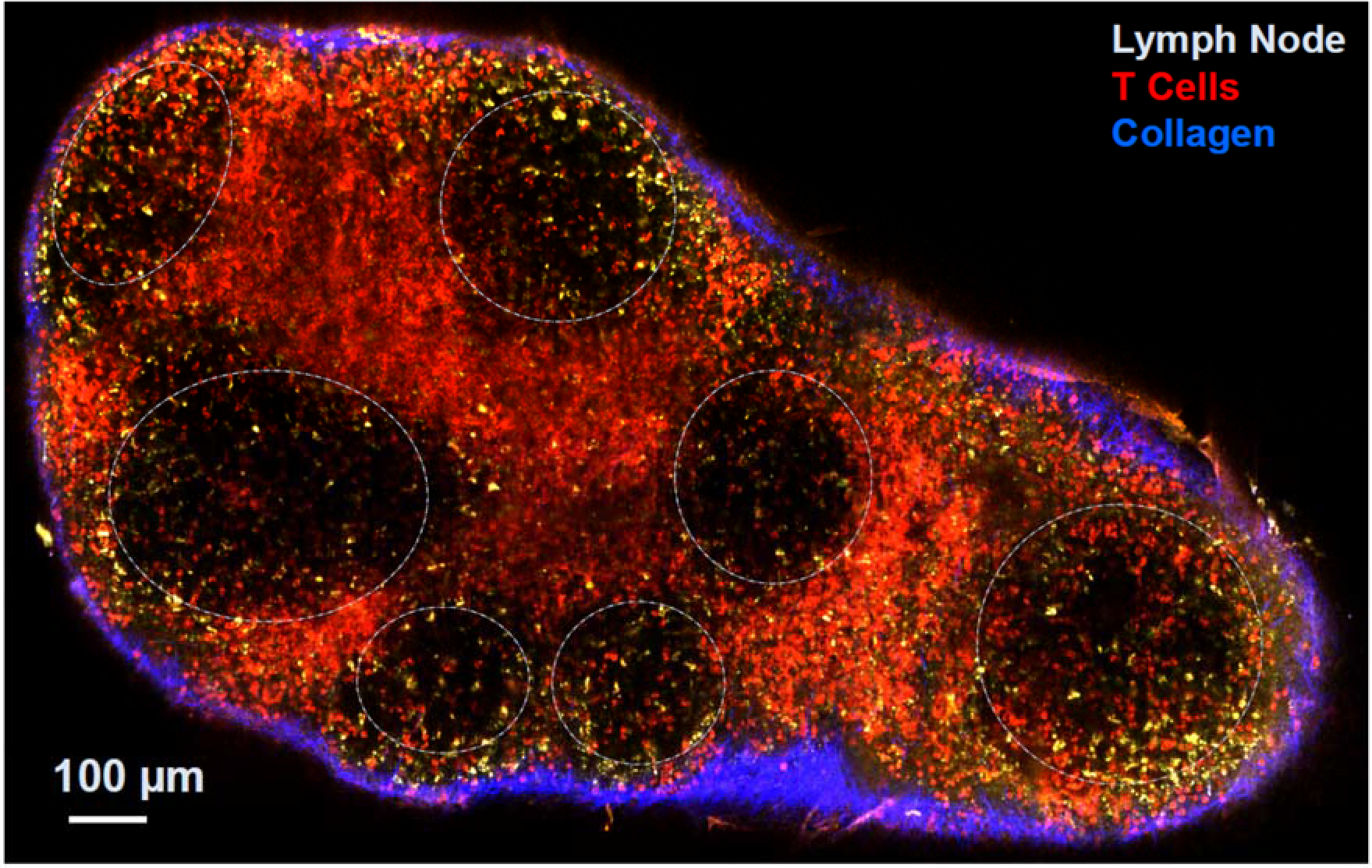

## Video Legends

**Video 1. Calcium readout of Salsa6f probe in HEK cells.** HEK 293A cells transfected with Salsa6f, first washed with 0 mM Ca^2+^ followed by 2 μM ionomycin in 2 mM Ca^2+^; scale bar = 20 μm, time shown in hr:min:sec. Images were acquired at 15 second interval and played back at 15 frames per second. This video corresponds to **Figure 1D**.

**Video 2. Single-cell readout of activation in transgenic T cells by Salsa6f**. CD4 T cells from CD4-Salsa6f^+/−^ mice were plated on activating surface coated with anti-CD3/CD28. Images were acquired at 5 second interval and played back at 15 frames per second. This video corresponds to **Figure 5A**.

**Video 3. T cell Ca^2+^ response to Ca^2+^ store depletion by thapsigargin (TG).** Video of maximum intensity projection images of 2 day activated T cells from CD4-Salsa6f^+/−^ mouse plated on poly-L-lysine. Scale bar = 20 μm, time shown in hr:min:sec. 2 μM TG in Ca^2+^ free Ringer’s was added at 00:02:30 and 2 mM Ca^2+^was added at 00:08:15. Time interval between frames is 5 sec. Play back speed = 50 frames per second. This video corresponds to **Figure 7A**.

**Video 4. Activated T cell Ca^2+^ responses to TCR stimulation.** Video of maximum intensity projection images of 2 day activated T cells from CD4-Salsa6f^+/−^ mouse plated on anti-CD3/28 coated coverslip. Scale bar = 20 μm, time shown in hr:min:sec. Time interval between frames is 5sec. Play back speed = 15 frames per second. Video corresponds to **Figure 7B**.

**Video 5. T cell Ca^2+^ response to shear and Yoda1.** Video of maximum intensity projection images of 2 day activated T cells from CD4-Salsa6f^+/−^ mouse plated on anti-CD3/28 coated coverslip. Scale bar = 20 μm, time shown in hr:min:sec. Time interval between frames is 5sec. Play back speed = 200 frames per second. Medium was added at 00:15:00and Yoda1 was added at 00:35:00. Video corresponds to **Figure 7C**.

**Video 6. Lymph nodes from CD4-Salsa6f^+/+^ mice exhibit cell-wide and subcellular Ca^2+^ signals.** Time shown in hr:min:sec; images were acquired at 5 second intervals. Play back speed = 50 frames per second. Red channel is turned off after beginning to facilitate visualization of green signals. Video corresponds to **Figure 9B**.

